# Contribution of innate and adaptive immune cells to the elimination of *Salmonella enterica* serotype Enteritidis infection in young broiler chickens

**DOI:** 10.1101/2021.01.29.428771

**Authors:** Nathalie Meijerink, Robin H.G.A. van den Biggelaar, Daphne A. van Haarlem, J. Arjan Stegeman, Victor P.M.G. Rutten, Christine A. Jansen

## Abstract

*Salmonella enterica* serotype Enteritidis (SE) is a zoonotic pathogen which causes foodborne diseases in humans through contaminated poultry products, as well as severe disease symptoms in young chickens. More insight in innate and adaptive immune responses of chickens to SE infection is needed to understand elimination of SE. Seven-day-old broiler chickens were experimentally challenged with SE and numbers and responsiveness of innate immune cells including natural killer (NK) cells, macrophages and dendritic cells (DCs) were assessed during 21 days post-infection (dpi). In parallel, numbers and function of γδ T cells, CD8^+^ and CD4^+^ T cells as well as antibody titres were determined. SE was observed in the intestine and spleen of SE-infected chickens at 7 dpi. NK and T cells responded first to SE at 1 and 3 dpi as indicated by increased numbers of intestinal IL-2Rα^+^ and 20E5^+^ NK cells, in addition to enhanced activation of intestinal and splenic NK cells. At 7 dpi in the spleen, the presence of macrophages and the expression of activation markers on DCs was increased. At 21 dpi, an increase in intestinal γδ and CD8^+^ T cell numbers was observed. Furthermore, SE-specific proliferation of splenic CD4^+^ and CD8^+^ T cells was observed and SE-specific antibodies were detected in all blood samples of SE-infected chickens. In conclusion, SE results in enhanced numbers and activation of innate cells during early stages of infection and it is hypothesized that in concert with subsequent specific T cell and antibody responses, reduction of SE in infected chickens is achieved. A better understanding of innate and adaptive immune responses important in the elimination of SE will aid in developing immune-modulation strategies, which may increase resistance and prevent SE infection and colonization in young broiler chickens and hence increase food safety for humans.

**Author summary:** *Salmonella enterica* serotype Enteritidis (SE) causes foodborne zoonotic diseases in humans, as well as a severe disease in young chickens. As a consequence of which health and welfare of humans and chickens are affected, resulting in substantial economic losses. To enable development of immune-mediated prevention strategies in chickens, more insight in the immune responses to SE is needed to understand how the infection is eliminated. For this purpose, we investigated non-specific and specific immune responses upon experimental SE infection in young broiler chickens. In this study, we found SE in the intestine and spleen of SE-infected chickens at 7 days post-infection (dpi). We show that natural killer (NK) cells respond first by enhanced presence and activation, followed by increased presence of macrophages and activation of dendritic cells. These early responses are hypothesized to stimulate the observed subsequent specific T cell and antibody responses. Better understanding of immune responses important in the elimination of SE will aid in developing immune-modulation strategies, which may increase resistance and prevent SE infection and colonization in young chickens and hence reduce SE-related foodborne illness in humans.

## Introduction

*Salmonella enterica* serotype Enteritidis (SE) is one of the leading causes of foodborne diseases in humans, most often due to poultry products as a prominent food source. In chickens infected with faecal *Salmonellae* via oral or respiratory routes, SE colonizes the intestinal tract and disseminate systemically to tissues such as the liver and spleen [1,2]. In young chickens it can lead to severe disease and death, whereas adult chickens are often subclinically infected with SE, carrying the bacteria in their intestines [3]. Prevention of SE infection in poultry is thus important for health and welfare of both young chickens and humans, as well as to avoid substantial economic losses as a consequence. Since therapeutic treatment of SE infection in chickens with antibiotics is not advised due to limited effectiveness and risk of antibiotic resistance, the use of immune-modulatory strategies to increase the resistance to SE is encouraged [4]. More insight in innate and adaptive immune responses and their interaction in response to SE infection in young broiler chickens will facilitate the design of these strategies. In young chickens, immunity largely depends on maternal antibodies as well activity of the innate immune system, with natural killer (NK) cells and macrophages as key players [5,6]. NK cells are particularly abundant amongst intestinal intraepithelial lymphocytes (IEL) [7,8], which are also rich in γδ T cells and CD8^+^ T cells expressing the αβ T cell receptor (TCR) [8,9]. Macrophages, dendritic cells (DCs) and CD4^+^ T cells are located directly underneath the intestinal epithelium [10]. The adaptive immune system is not fully developed yet upon hatch and functional T- and B- cell responses are observed after approximately two to three weeks of life [5,6].

The early response of the innate immune system in chickens within one week post-SE infection is characterised by the upregulation of genes associated with ‘defense/pathogen response’ [11], NK cell-mediated cytotoxicity [12] and production and secretion of the cytokine IFNγ [13]. In chickens, intestinal NK cells comprise a major CD3^−^ IL-2Rα^+^ subset [7,8] and a minor CD3^−^ 20E5^+^ subset [7], both having cytotoxic capacity but to different degrees. In mice and humans, high cytotoxic activity [14,15] and IFN-γ production [15,16] by NK cells have been shown to result in resistance to *Salmonella enterica* serotype Typhimurium. In chickens, the effect of SE infection on the function of NK cells has not been studied so far.

The development of T and B cell responses is initiated with the activation of antigen presenting cells (APCs) such as DCs and macrophages. The presence [17] and activity of intestinal macrophages [18–20], identified by the expression of mannose receptor C-type 1-like B (MRC1LB) [21], and bone marrow-derived DCs [22] increases during SE infection. Furthermore, oral infection of 1-day-old specific-pathogen-free chickens with SE elicits increased mRNA expression of chemokines and macrophages are attracted to the ileum within 24 hours [23]. In addition, it has been shown that decreased activity of peritoneal macrophage is associated with increased susceptibility for systemic dissemination of SE in chickens [24]. On the other hand, *Salmonella* species have been found to resist killing by macrophages in mammals [25] and chickens [26,27] and even use macrophages as a carrier for systemic dissemination [28]. Despite their involvement in SE infection, a detailed analysis of the effect of SE infection on the function of APCs in chickens has not been performed to date.

Initial T cell responsiveness to SE in chickens has been observed within one week, including increased presence of γδ T cells in the intestine, blood and spleen after live vaccination [13,29], and enhanced mRNA expression of cytotoxic activity-related genes in the spleen after infection [30], as compared to non-vaccinated and uninfected chickens. After two and three weeks, a second increase in presence of γδ T cells was observed in the spleen and intestine, respectively, compared to non-vaccinated chickens [29]. CD4^+^ helper T cells and CD8^+^ cytotoxic T cells were also shown to increase in presence within one week post-SE infection and vaccination in the intestine, as compared to uninfected and non-vaccinated chickens [13,17,18]. CD8^+^ cytotoxic T cells kill infected host cells, while CD4^+^ helper T cells release cytokines like IL-2 and IFNγ to further stimulate NK cells, and macrophages and CD8^+^ cytotoxic T cells respectively, and promote the differentiation of B cells into antibody-producing plasma cells. Antibody responses involved in elimination of SE partly depend on maternal antibodies and on the production of IgA in the intestine [31,32], as well as IgM, IgA and IgY antibodies in blood [32,33]. Whereas previous studies have focussed on specific aspects of the immune responses, the present study combines cellular assays to analyze both the innate and adaptive immune responses in intestine and spleen upon SE infection in chickens.

In this study, we investigated how *Salmonella enterica* serotype Enteritidis infection in young broiler chickens affects presence and activation of innate and adaptive immune cells in the intestine and spleen to obtain more insight in the contribution of the immune system to elimination of the infection. Numbers of SE in ileum and spleen were determined alongside differences in kinetics of presence and activation status of NK cell and APC subsets between uninfected and infected chickens. The subsequent adaptive responses were determined including presence and activation of γδ T cell, CD4^+^ and CD8^+^ T cell subsets, and serum antibody levels. Hence the present study provides an extensive overview of intestinal and systemic immune responses that are evoked by SE infection in young broiler chickens. Based on the phenotypical and functional data obtained, we will hypothesize on how the various elements of the immune system interact and contribute to elimination of the SE infection, and on potential strengthening of immune responsiveness by immunomodulation strategies, which may prevent SE infection and colonization, and thus increase chicken health and welfare as well as safety of food of chicken origin.

## Materials and methods

### Animals and husbandry

A total of 30 respectively 35 Ross 308 seventeen- and eighteen-day old embryonated eggs were obtained from the same parent flock of a commercial hatchery (Lagerwey, the Netherlands). Eggs were disinfected with 3% hydrogen peroxide and placed in disinfected hatchers in two different stables (uninfected and SE-infected) at the facilities of the Department of Population Health Sciences, Faculty of Veterinary Medicine, Utrecht University, the Netherlands. Directly upon hatch, chickens were weighed, labelled and housed in floor pens lined with wood shavings (2 kg/m^2^), and received water and standard *Salmonella*-free commercial starter and grower feeds *ad libitum* (Research Diet Services, the Netherlands). A standard lighting and temperature scheme for Ross broiler chickens was used for both stables.

The animal experiment was approved by the Dutch Central Authority for Scientific Procedures on Animals and the Animal Experiments Committee (registration number AVD1080020174425) of Utrecht University (the Netherlands) and all procedures were done in full compliance with all relevant legislation.

### Experimental design

Before the start of the experiment at day 3, five chickens per group (uninfected (*n* = 30) and SE-infected (*n* = 35)) were randomly selected and sacrificed for collection of ileum (±10 cm distal from Meckel’s diverticulum) and spleen to confirm absence of SE before inoculation. At day 7 (0 days post infection (dpi)), five chickens of the SE-infected group only, were randomly selected and sacrificed for collection of ileum and spleen, to determine baseline levels of the various immune parameters as well as absence of SE before infection. Subsequently, chickens of the SE-infected group were challenged at day 7 (0 dpi) by oral inoculation of 0.25 ml brain heart infusion (BHI) medium containing 1.12×10^6^ colony-forming units (CFUs) SE, whereas chickens in the other stable (uninfected) were inoculated with 0.25 ml BHI medium. At days 8 (1 dpi), 10 (3 dpi), 14 (7 dpi), 21 (14 dpi) and 28 (21 dpi), five chickens per group were randomly selected and sacrificed for collection of ileum and spleen to determine bacterial CFUs as well as numbers and function of NK cells and T cells. At days 7, 8, 10 and 14 also spleen APCs were assessed. At days 7, 14, 21 and 28 blood (at least 5 ml) was collected in EDTA tubes (VACUETTE^®^ K3E EDTA, Greiner Bio-One, the Netherlands) for determination of SE-specific antibody levels. At day 28, splenic lymphocytes were also used to assess SE-specific T cell reactivity in a proliferation assay. All chickens were weighed prior to post-mortem analyses to determine the growth curve. Ileum segments and spleens were weighed before cell isolation to enable calculation of absolute cell numbers.

### SE culture

The *Salmonella enterica* serotype Enteritidis strain (K285/93 Nal^res^) was kindly provided by Dr. E. Broens, director of the Veterinary Microbiological Diagnostic Center (VMDC) of the Faculty of Veterinary Medicine, Utrecht University, and cultured as described previously [34]. In short, from an overnight culture of the SE strain on blood agar (Oxoid, the Netherlands) a single colony was used to inoculate 45 ml BHI medium (Oxoid), which was incubated aerobically overnight at 200 rpm in a shaking incubator at 37°C. The OD value of a sample of the SE culture diluted 1:10 in PBS was measured using a Ultrospec 2000 (Pharmacia Biotech, Sweden), the SE concentration was calculated from a previously determined growth curve, and SE were diluted in BHI medium to 4 × 10^6^ CFU/ml, to constitute the inoculum. The exact SE concentration of the inoculum, determined by counting the number of CFUs of plated serial dilutions after overnight culture, was 4.49 × 10^6^ CFU/ml.

For the T cell proliferation assay, SE was fixed by resuspending 3.8×10^9^ CFUs in 100 μl PBS with 1% formaldehyde (Sigma-Aldrich, the Netherlands) and incubation for 5 min at RT, while the suspension was vortexed shortly every minute. After fixation, the bacteria were washed four times in 1 ml PBS by centrifugation at 12500 rpm to remove the supernatants (Heraeus Pico 17 Centrifuge, Thermo Fisher Scientific). Finally, the bacteria were resuspended in 380 μl X-VIVO 15 cell culture medium (Lonza, the Netherlands) with 50 μg/ml gentamycin (Gibco™, the Netherlands) to create a concentration of 10^7^ CFU/ml and stored at 4°C until further use.

### Isolation of cells

Ileum segments were washed with PBS to remove the contents, cut into sections of 1 cm^2^ and washed again. Subsequently, the IEL were collected by incubating three times in a shaking incubator at 200 rpm for 15 min at 37°C in EDTA-medium (HBSS 1x (Gibco^®^) supplemented with 10% heat-inactivated FCS (Lonza) and 1% 0.5M EDTA (Sigma-Aldrich). Supernatants were collected and centrifuged for 5 min at 1200 rpm at 20°C (Allegra™ X-12R Centrifuge, Beckman Coulter, the Netherlands). Cells were then resuspended in PBS, lymphocytes were isolated using Ficoll-Paque Plus (GE Healthcare, the Netherlands) density gradient centrifugation for 12 min at 1700 rpm at 20°C, washed in PBS by centrifugation for 5 min at 1300 rpm at 4°C and resuspended at 4.0 × 10^6^ cells/ml in complete medium (IMDM 2 mM glutamax I supplemented with 8% heat-inactivated FCS (Lonza), 2% heat-inactivated chicken serum, 100 U/ml penicillin/streptomycin; Gibco^®^). Spleens were homogenized using a 70 μm cell strainer (Beckton Dickinson (BD) Biosciences, NJ, USA) to obtain a single-cell suspension. Next, lymphocytes were isolated by Ficoll-Paque density gradient centrifugation (20 min, 2200 rpm, 20°C), washed in PBS and resuspended at 4.0 × 10^6^ cells/ml in complete medium as described for ileum.

Whole blood was allowed to coagulate by leaving it undisturbed for 1 hour at room temperature (RT), centrifuged for 10 min at 3000 rpm at 15°C and subsequently, serum was collected and stored at −20°C until further use.

### Quantitative bacteriology of IEL and spleen

At −4, 0, 1, 3, 7, 14 and 21 dpi, the numbers of *Salmonella* colonies in IEL and spleen samples were determined by plating 100 μl of the EDTA-isolated IEL and homogenized spleen, 1:10-diluted in PBS before the Ficoll step, on RAPID’*Salmonella* Medium plates (Bio-Rad, the Netherlands). Plates were incubated overnight at 37°C and subsequently, purple colonies were quantified and SE was expressed as CFU per gram tissue.

### Phenotypic characterization of lymphocytes by flow cytometry

Presence and activation of NK and T cell subsets were determined among IEL and splenocytes at 0, 1, 3, 7, 14 and 21 dpi as described previously [7,35]. Lymphocyte populations (1 × 10^6^) were stained with a panel of antibodies specific for surface markers known to be expressed on NK cells, as well as with anti-CD3 to exclude T cells from the analyses. In addition, cells were stained with a panel of antibodies specific for surface markers that distinguishes γδ T cell, CD4^+^ and CD8^+^ T cell subsets (Table 1). Staining with primary and secondary antibodies was performed in 50 μl PBS (Lonza) containing 0.5% bovine serum albumin and 0.1% sodium azide (PBA). Cells were incubated for 20 min at 4°C in the dark, washed twice by centrifugation for 5 min at 1300 rpm at 4°C in PBA, after primary staining, and in PBS after secondary staining. Subsequently, to be able to exclude dead cells from analysis, lymphocytes were stained in 100 μl PBS with a viability dye (Zombie Aqua™ Fixable Viability Kit, Biolegend, the Netherlands) for 15 min at RT in the dark, washed twice in PBA and resuspended in 200 μl PBA. Of each sample, either 150 μl or a maximum of 1 × 10^6^ viable lymphocytes were analyzed using a CytoFLEX LX Flow Cytometer (Beckman Coulter), and data was analyzed with FlowJo software (FlowJo LCC, BD Biosciences). The gating strategies used to analyze NK cells, γδ T cells and cytotoxic CD8^+^ T cells were as described previously [35]. In short, gating included consecutive selection for lymphocytes (FSC-A vs SSC-A), viable cells (Live/Dead marker-negative) followed by selection of the cell subsets and activation markers according to the staining panels. NK cell subsets: CD3^−^ cells expressing either IL-2Rα or 20E5. NK activation: CD3^−^CD41/61^−^ cells expressing CD107 or CD3^−^ cells expressing IFNγ. T cell subsets: CD3^+^CD4^−^ cells positive for TCRγδ (γδ) or negative (CD8^+^ αβ) with both γδ and cytotoxic αβ T cells expressing either CD8αα or CD8αβ, and only in spleen the CD3^+^ cells expressing CD4. T cell activation: CD3^+^CD41/61^−^CD8α^+^ cells expressing CD107 or CD3^+^TCRγδ^+^CD8α^+^, CD3^+^TCRγδ^−^CD8α^−^ (CD4^+^) and CD3^+^TCRγδ^−^CD8α^+^ cells expressing IFNγ.

**Table 1.** Flow cytometry staining reagents.

| Cell population | Primary antibody (mouse-anti-chicken) | Clone / Isotype | Secondary antibody |
| --- | --- | --- | --- |
| NK cells | CD45-FITC<sup>1</sup> | LT40 / IgM | - |
|  | CD3-APC<sup>1</sup> | CT3 / IgG1 | - |
| | IL-2Rα-UNL<sup>2</sup> | 28-4 / IgG3 | Goat-anti-mouse-IgG3-PE<sup>1</sup> |
|  | 20E5-BIOT<sup>3</sup> | IgG1 | Streptavidin (SA)-PercP<sup>6</sup> |
| T cells | CD3-PE<sup>1</sup> | CT3 / IgG1 | - |
|  | CD4-APC<sup>1</sup> | CT4 / IgG1 | - |
| | TCR$\gamma\delta$-FITC<sup>1</sup> | TCR-1 / IgG1 | - |
| | CD8α-UNL<sup>1</sup> | EP72 / IgG2b | Goat-anti-mouse-IgG2b-APC/Cy7<sup>1</sup> |
| | CD8β-BIOT<sup>1</sup> | EP42 / IgG2a | SA-PercP<sup>6</sup> |
| APCs | CD41/61-FITC<sup>4</sup> | 11C3 / IgG1 | - |
|  | Bu-1-AF647<sup>1</sup> | AV20 / IgG1 | - |
|  | CD3-FITC<sup>1</sup> | CT3 / IgG1 | - |
|  | CD4-PE/Cy7<sup>1</sup> | CT4 / IgG1 | - |
|  | MRC1LB-PE<sup>1</sup> | KUL01 / IgG1 | - |
|  | CD11-biotin<sup>3</sup> | 5C7 / IgG1 | SA-Brilliant Violet (BV) 605<sup>7</sup> |
|  | MHC-II-UNL<sup>1</sup> | Cia / IgM | Rat-anti-mouse-IgM-BV421 (RMM-1)<sup>7</sup> |
|  | *CHIR-AB1-UNL<sup>2</sup> | 8D12 / IgG2a | Rat-anti-mouse-IgG2a-PerCP/Cy5.5 (RMG2a-62)<sup>7</sup> |
|  | *CD40-UNL<sup>5</sup> | AV79 / IgG2a | Rat-anti-mouse-IgG2a-PerCP/Cy5.5 (RMG2a-62)<sup>7</sup> |
|  | *CD80-UNL<sup>5</sup> | IAH:F864:DC7 / IgG2a | Rat-anti-mouse-IgG2a-PerCP/Cy5.5 (RMG2a-62)<sup>7</sup> |
| **Assay** |  |  |  |
| CD107 | CD107a-APC<sup>3</sup> | LEP-100 I 5G10 / IgG1 | - |
|  | CD41/61-FITC<sup>4</sup> | 11C3 / IgG1 | - |
|  | CD3-PE<sup>1</sup> | CT3 / IgG1 | - |
| | CD8α-UNL<sup>1</sup> | EP72 / IgG2b | Goat-anti-mouse-IgG2b-Alexa Fluor (AF) 790<sup>8</sup> |
|  | 28-4-UNL<sup>2</sup> | IgG3 | Goat-anti-mouse-IgG3-APC/Cy7<sup>1</sup> |
|  | 20E5-BIOT<sup>3</sup> | IgG1 | SA-PercP<sup>6</sup> |
| IFNγ | CD3-PE<sup>1</sup> | CT3 / IgG1 | - |
| | TCR$\gamma\delta$-FITC<sup>1</sup> | TCR-1 / IgG1 | - |
| | CD8α-UNL<sup>1</sup> | EP72 / IgG2b | Goat-anti-mouse-IgG2b-AF790<sup>8</sup> |
|  | 28-4-UNL<sup>2</sup> | IgG3 | Goat-anti-mouse-IgG3-APC/Cy7<sup>1</sup> |
|  | 20E5-BIOT<sup>3</sup> | IgG1 | SA-PercP<sup>6</sup> |
| | IFNγ-APC<sup>3</sup> | MAb80 / IgG1 | - |
| T cell proliferation | CD3-FITC<sup>1</sup> | CT3 / IgG1 | - |
|  | CD4-PE/Cy7<sup>1</sup> | CT4 / IgG1 | - |
| | CD8α-APC<sup>1</sup> | CT8 / IgG1 | - |
| | IL-2Rα-UNL<sup>2</sup> | 28-4 / IgG3 | Goat-anti-mouse-IgG3-PE<sup>1</sup> |

Three APC antibody panels were prepared each containing surface markers plus one out of three antibodies indicated (\*). Manufacturer: <sup>1</sup>Southern Biotech, AL, USA, <sup>2</sup>Purified supernatant of hybridoma provided by Göbel, T.W., Ludwig Maximilian University, Germany, <sup>3</sup>Developmental Studies Hybridoma Bank (DSHB), University of Iowa, IA, USA, <sup>4</sup>Serotec, United Kingdom, <sup>5</sup>Bio-Rad, <sup>6</sup>BD Biosciences, <sup>7</sup>Biolegend, <sup>8</sup>Jackson ImmunoResearch Laboratories, PA, USA.

### CD107 assay

Activation of NK cells and cytotoxic CD8^+^ T cells was determined using the CD107 assay, which measures the increased surface expression of CD107a that results from degranulation of cytotoxic granules [36]. Briefly, lymphocytes isolated from IEL and spleen were resuspended in complete medium, and 1 × 10^6^ lymphocytes in 0.5 ml were incubated in the presence of 1 μl/ml GolgiStop (BD Biosciences) and 0.5 μl/ml mouse-anti-chicken-CD107a-APC for 4 hours at 37°C, 5% CO_2_. After incubation, lymphocytes were washed in PBA and stained as described in 2.6 with monoclonal antibodies for NK and T cells, and anti-CD41/61 to exclude thrombocytes from analyses, as mentioned in the CD107 panel (Table 1). Cells were washed in PBS, stained for viability and analyzed by flow cytometry as described in 2.6.

### IFNγ assay

Expression of intracellular IFNγ was determined in (subsets of) NK cells, γδ T cells, CD4^+^ and CD8^+^ T cells, using the assay adapted from Ariaans et al. 2008 [37]. Lymphocytes isolated from IEL and spleen were resuspended in complete medium, and 1 × 10^6^ lymphocytes in 0.5 ml were incubated in the presence of 1 μl/ml Brefeldin A (Sigma Aldrich) for 4 hours at 41°C, 5% CO_2_. After incubation, lymphocytes were washed in PBA and stained as described in 2.6 with surface markers as mentioned in the IFNγ panel (Table 1). Cells were washed in PBS, stained for viability and washed again in PBA. Then, lymphocytes were permeabilized differently by using different solutions as described before. Lymphocytes were incubated in 200 μl of a mixture of BD FACS™ Permeabilizing Solution 2 and BD FACS™ Lysing Solution prepared according to manufacturer’s instructions (BD Biosciences) for 8 min at RT, immediately followed by centrifugation for 2 min at 1300 rpm at 4°C. Cells were washed twice in PBA, stained intracellularly with anti-IFNγ-APC in 50 μl PBA for 20 min at 4°C in the dark, washed in PBA and finally analyzed by flow cytometry as described in 2.6.

### Phenotypic characterization of APCs by flow cytometry

Splenocytes isolated at 0, 1, 3 and 7 dpi from infected and uninfected chickens were transferred to a 96 wells V-bottom plate and stained with antibodies of the APC panel to distinguish APC subsets (Table 1). Staining with primary and secondary antibodies (1 × 10^6^) was performed in 50 μl PBA, incubated for 20 min at 4°C in the dark and washed twice by centrifugation for 3 min at 1300 rpm at 4°C in PBA. Finally, the cells were stained in 50 μl PBS with ViaKrome 808 viability dye (Beckman Coulter) for 20 min at 4°C. Cells were washed in PBA and analyzed by flow cytometry as described in 2.6, using 180 μl.

Based on the APC subset staining, a t-distributed Stochastic Neighbor Embedding (t-SNE) analysis was performed using FlowJo software to identify cell subsets using an unbiased approach. From each sample of splenocytes at 7 dpi of infected (*n* = 5) and uninfected (*n* = 5) chickens, 10,000 cells were taken and concatenated into one FCS file that represented all individual chickens. The t-SNE was performed based on expression levels of CD3, CD41/61, MRC1LB, CD4, Bu-1, CD11, MHC-II and CHIR-AB1 using published automated optimized parameters [38]. Based on the t-SNE, three APC subsets were identified based on selection of MRC1LB and CD11 positive cells, which were negative for CD3, CD4 and CD41/61. Activation status of the subsets was subsequently evaluated using the expression percentages of immunoglobulin Y receptor CHIR-AB1, co-stimulatory molecules CD40 and CD80, and the geometric mean fluorescent intensity (gMFI) of MHC-II. CHIR-AB1 was included as an activation marker since van den Biggelaar et al. (2020) recently showed increase of its expression on macrophages after stimulation with LPS [39].

### Fluorescence-activated cell sorting of NK cell and APC subsets

Based on marker expression, two NK cell subsets and three APC subsets were separated by fluorescence-activated cell sorting (FACS) to gain more insight into their functional identity. To distinguish NK cell subsets, splenocytes were stained with mouse-anti-chicken-CD3-APC, -Bu-1-FITC, −28-4 and −20E5-biotin. For secondary antibody staining goat-anti-mouse-IgG3-PE and SA-BV421 were used. To identify APC subsets, splenocytes were stained with mouse-anti-chicken-MRC1LB-PE and -CD11-biotin. Secondary staining with SA-BV605 was used to fluorescently label CD11-biotin. To assess viability, the cells were stained with the Zombie Aqua™ Fixable Viability Kit. Primary and secondary antibody staining of cells used the same conditions as described in 2.6. Finally, the cells were resuspended in PBA (NK cells) or PBA with 2 mM EDTA (APC subsets), and isolated by FACS using the BD influx™ Cell Sorter and 405-, 488-, 638-, 561- and 640-nm lasers. The cells were gated for viability and subsequently sorted into CD3^−^ Bu-1^−^ 28-4^+^ (IL-2Rα^+^ NK), CD3^−^ Bu-1^−^ 20E5^+^ (20E5^+^ NK), respectively CD11^+^ MRC1LB^+^ (APC subset 1), CD11^+^ MRC1LB^−^ FSC^low^ (APC subset 2a) and CD11^+^ MRC1LB^−^ FSC^high^ (APC subset 2b). The sorted NK cell subsets were collected in 350 μl RLT buffer (Qiagen, the Netherlands) with 1% 2-mercaptoethanol (Sigma Aldrich). The sorted APC subsets, and the original (unsorted) cell population as a control, were centrifuged at 1300 rpm for 5 min and then lysed in 600 μl RLT buffer with 1% 2-mercaptoethanol. Cell lysates of sorted cell subsets and the control cell population were then stored at −20°C until RNA isolation and qPCR analysis.

### Gene expression of separated subsets of NK cells and APCs

Target genes (Table 2) were selected based on literature to define functional differences between subsets of NK cells respectively APCs. RNA was isolated from lysates of sorted NK and APC subsets, and control cells, using the RNeasy Mini Kit (Qiagen) according to the manufacturer’s instructions, including a DNase treatment using the RNase-Free DNase Set (Qiagen). Next, cDNA was prepared using the reverse transcriptase from the iScript cDNA Synthesis Kit (Bio-Rad) according to the manufacturer’s instructions. RT-qPCRs were performed with primers and either FAM-TAMRA-labeled TaqMan probes combined with TaqMan Universal PCR Master Mix or SYBR Green Master Mix without probes (all from Thermo Fisher Scientific), as indicated in Table 2. Primers were used at 400 nM (SYBR-Green) or 600 nM (Taqman) and probes at 100 nM. RT-qPCRs were performed with a CFX Connect and analyzed with CFX Maestro software (both from Bio-Rad). All RT-qPCRs were evaluated for proper amplification efficiency (95–105%) using serial dilutions of reference cDNA either from splenocytes that were stimulated with Concanavalin A for 24 hours or from HD11 cells that were stimulated with LPS for 3 hours. RT-qPCRs were performed in triplicate for every sample. For the NK cell subsets, mRNA levels are expressed as 40-Ct and the cycle threshold value (Ct) was corrected for variations in RNA preparation and sampling using the *GAPDH* Ct values, as described elsewhere [40]. Higher gene expression of *NFIL3* and *IL-7α* is indicative of the cytokine-producing NK cell subset in humans [41–45], whereas higher expression of *PRF1* is indicative of the cytotoxic NK cell subset in humans and chickens [43,46]. Gene expression levels of the APC subsets are shown relative to those of unsorted splenocytes. Furthermore, Ct gene expression values were normalized to housekeeping genes *28S* and *GAPDH*. Changes in gene expression after sorting was expressed as 2^−ΔΔCt^, according to the Livak method [47]. Enrichment of cells expressing *CD14*, *TLR4*, *MERTK* and *MAFB* after sorting of cells was considered indicative for a monocyte/macrophage phenotype, whereas enrichment of cells expressing *ZBTB46*, *XCR1* and *FLT3* after sorting was considered indicative for a DC phenotype, in accordance with previous studies [48,49].

**Table 2.**
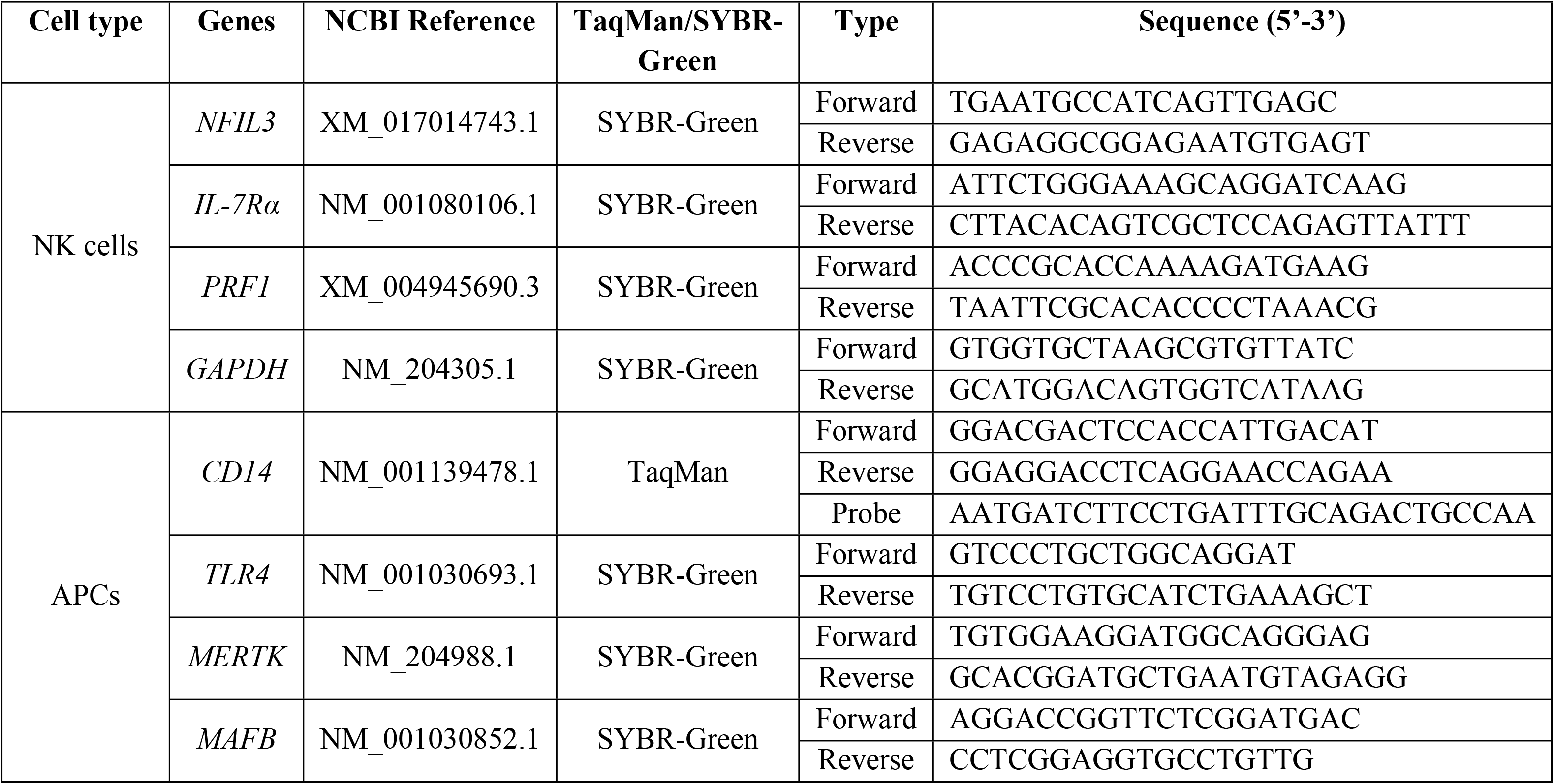
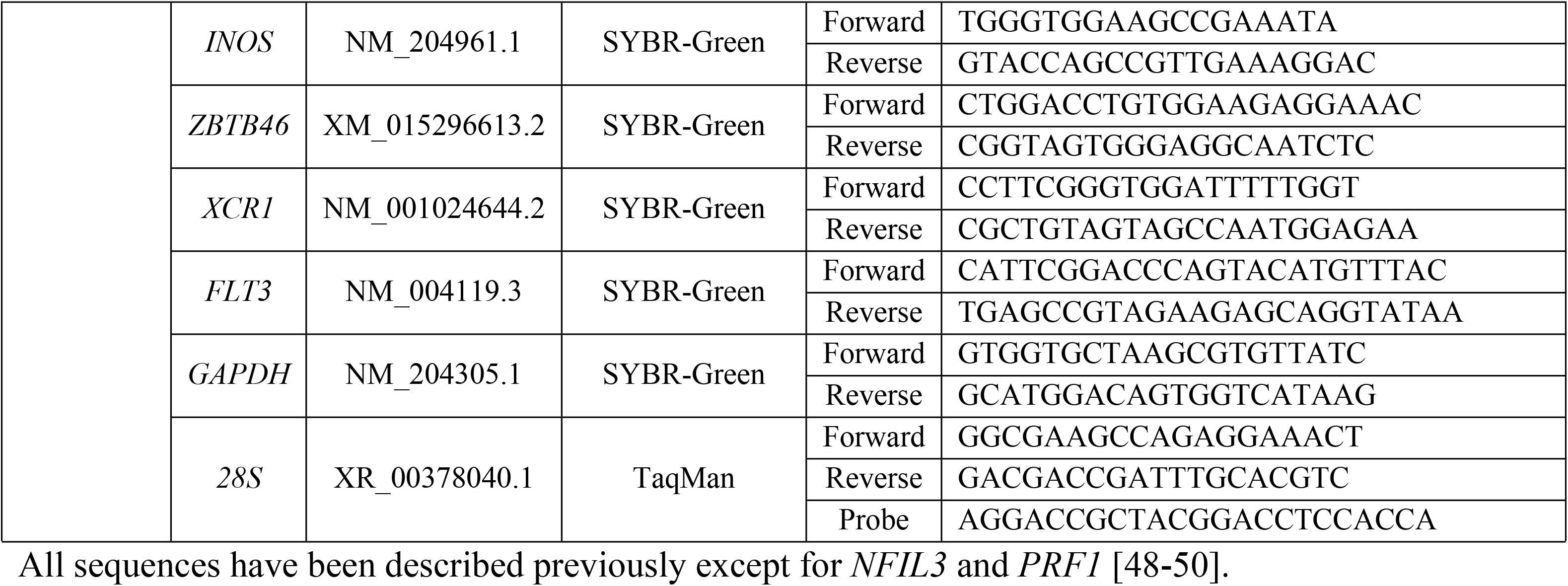
Primers and TaqMan probe sequences used for RT-qPCR.

### T cell proliferation assay

Splenocytes isolated at 21 dpi from uninfected and SE-infected chickens were labelled with CellTrace Violet (CTV, Invitrogen, the Netherlands) to measure proliferation by flow cytometry. The cells were resuspended at 5×10^6^ cells/ml in PBS with 5 μM CTV and incubated for 20 min at RT, while the cell suspension was vortexed every 5 min. Next, the labeling was quenched by the addition of 5 ml complete medium for every ml of CTV staining solution and incubated for 5 min at RT. Cells were centrifuged for 5 min at 1200 rpm at 20°C and resuspended at 2.5×10^6^ cells/ml in X-VIVO 15 cell culture medium (Lonza) with 50 U/ml penicillin–streptomycin, 50 μM 2-mercaptoethanol (Sigma Aldrich) and 50 μg/ml gentamycin (Gibco™). Aliquots of 200 μl cell suspension containing 500,000 splenocytes were added to the wells of a 96 wells round-bottom cell culture plate. Fixed SE was added to the splenocytes at 10^4^, 10^5^ or 10^6^ CFU/well. As a positive control, splenocytes were stimulated with 1 μg/ml mouse-anti-chicken-CD3, 1 μg/ml -CD28 and 1:50 diluted conditioned supernatant from COS-7 cells transfected with a pcDNA1 vector (Invitrogen) encoding for recombinant chicken IL-2 (a kind gift from prof. Pete Kaiser and dr. Lisa Rothwell), in accordance with a previous publication [51]. Cells were incubated for four days at 41°C and 5% CO_2_. After incubation, cells were transferred to a 96 wells V-bottom plate and stained in PBS with ViaKrome 808 viability dye (Beckman Coulter). Next, cells were stained with antibodies of the T cell proliferation panel (Table 1). Primary and secondary staining of cells were conducted in 30 μl PBA and incubated for 20 min at 4°C in the dark. Stained cells were washed twice by centrifugation for 3 min at 1300 rpm at 4°C in PBA and resuspended in 100 μl followed by flow cytometry analysis as described in 2.6, using 80 μl.

### SE-specific antibody titers in serum

To detect titers of SE-specific antibodies in the sera collected at 0, 7, 14 and 21 dpi, the commercially available *Salmonella* Enteritidis Antibody Test (IDEXX SE Ab X2 Test) was used according to manufacturer’s instructions (IDEXX Europe, the Netherlands). Positive and negative controls were included in the kit, and serum samples were analyzed in duplicate. Endpoint titers were calculated with the following formula:

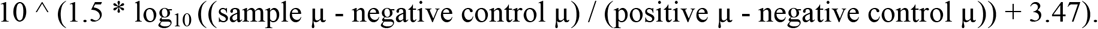

### Statistical analysis

First, the data were tested for fitting a normal distribution using the Shapiro-Wilk test. Differences in numbers of leukocytes, NK cell and T cell subsets and percentages of CD107 and IFNγ expression in IEL and spleen between the uninfected and SE-infected groups were analyzed using one-way ANOVA tests. Differences in SE CFUs per gram intestine and spleen as well as SE-specific antibody titers in serum were analyzed using Kruskal-Wallis tests accompanied by Dunn’s multiple comparisons tests. Differences in numbers and percentages of the splenic APC subsets 1 and 2a between the uninfected and SE-infected groups were analyzed using one-way ANOVA tests, while subset 2b was analyzed using the Kruskal-Wallis test as the data was not normally distributed. All statistical analyses were performed using GraphPad Prism 8 software (GraphPad Software, CA, USA). A *p*-value of < 0.05 was considered statistically significant.

## Results

### Highest presence of SE in ileum and spleen at 7 dpi while intestinal infiltration of leukocytes was observed at 1 and 3 dpi

SE was not observed at −4 and 0 dpi before SE inoculation, in both the IEL fraction of the ileum and the spleen of chickens of both groups (Fig 1A and 1B). After inoculation SE was detected in the IEL fraction of SE-infected chickens only at 7 dpi (11,100±4026 CFU/g, mean±SEM, Fig 1A). In the spleen, SE was observed at 7, 14 and 21 dpi with the highest bacterial counts at 7 dpi (1200±566 CFU/g), which subsequently decreased in course of time (Fig 1B). SE was not detected in the IEL fraction and spleen of uninfected chickens at any of the time points (Fig 1A and 1B). One uninfected chicken showed counts of *Proteus* in the spleen at 7 dpi and was therefore excluded from further analyses. Infection with SE did not affect the weight of the chickens, as growth curves were similar between uninfected and SE-infected chickens (Fig 1C). A significant increase in numbers of intestinal leukocytes was found in SE-infected chickens at 1 dpi (38,509±3544) compared to uninfected chickens (15,401±1716, Fig 1D). Numbers tended to be higher at 3 dpi in SE-infected chickens and declined over time to numbers similar to those observed in uninfected chickens (Fig 1D). The numbers of splenic leukocytes were similar between uninfected and SE-infected chickens at all time points (Fig 1E).

**Fig 1.**
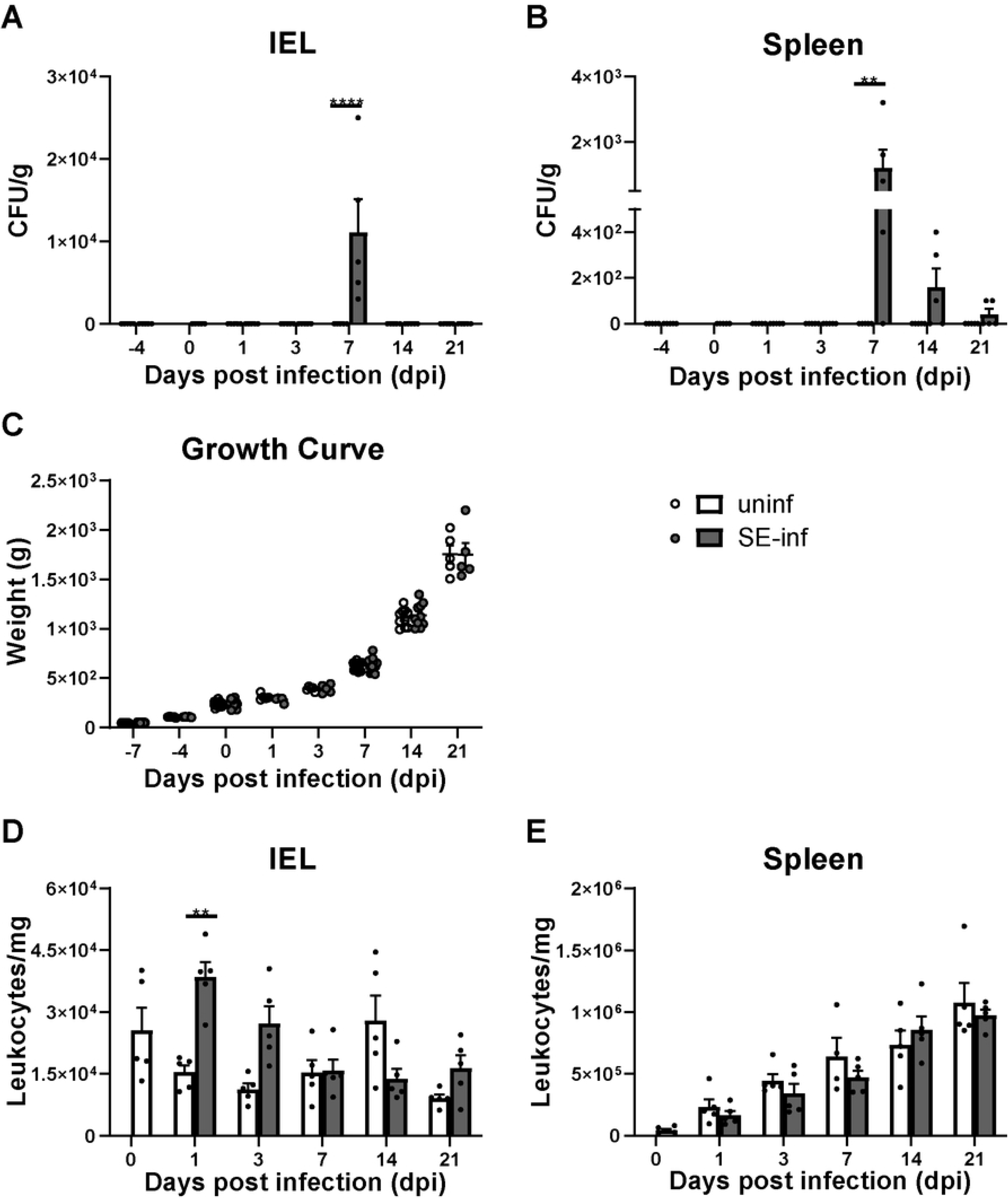
The bacterial load in course of time after SE infection and its effect on growth and numbers of intestinal and splenic leukocytes. (A) *Salmonella enterica* serotype Enteritidis (CFU/g) in the IEL fraction of the ileum and in the (B) spleen of uninfected (uninf) and SE-infected chickens (SE-inf). (C) Bodyweights (g) of uninfected and SE-infected chickens in the course of time. (D) Numbers (cells/mg) of leukocytes in IEL and (E) spleen of uninfected and SE-infected chickens in the course of time. Mean + SEM per treatment and time point is shown (*n* = 5) and statistical significance is indicated as ** *p* < 0.01 and **** *p* < 0.0001.

### Increased presence and enhanced activation of intestinal NK cells upon SE infection

Numbers of intestinal and splenic NK cell subsets were determined to investigate differences between uninfected and SE-infected chickens at several time points post-infection. NK cell subsets were distinguished by membrane expression of IL-2Rα or 20E5 (Fig 2A and S1A Fig). In the intestine, numbers of IL-2Rα^+^ and 20E5^+^ NK cells tended to be higher in SE-infected chickens at 1, 3 and 7 dpi compared to uninfected chickens (Fig 2B and 2C). Intestinal 20E5^+^ NK cell numbers tended to be lower at 14 dpi and higher at 21 dpi in SE-infected compared to uninfected chickens (Fig 2C). In the spleen, numbers of IL-2Rα^+^ and 20E5^+^ NK cells were similar between uninfected and SE-infected chickens and both increased in course of time (S1B and S1C Fig). To obtain more insight in functional differences between IL-2Rα^+^ and 20E5^+^ NK cells, mRNA levels of genes deemed to be relevant were determined in the spleen. The IL-2Rα^+^ subset showed higher mRNA levels of the NK cell lineage marker NFIL3 and IL-7Rα^+^ as compared to the 20E5^+^ subset, whereas the 20E5^+^ subset showed higher mRNA levels of perforin as compared to the IL-2Rα^+^ subset (PRF1, S1D Fig).

**Fig 2.**
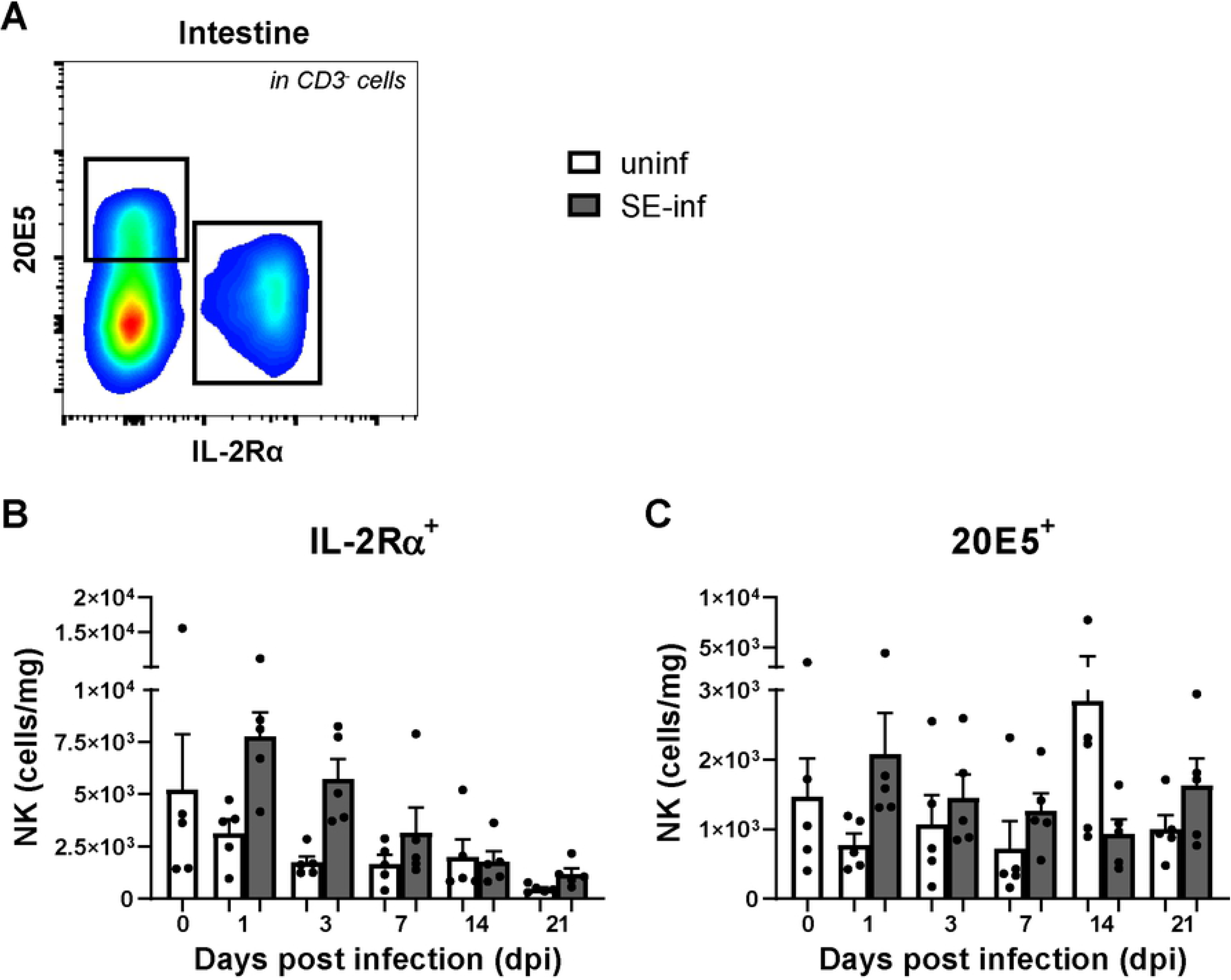
Effect of SE infection on numbers of intestinal NK cells in broiler chickens. (A) Gating strategy for CD3 negative cells expressing either IL-2Rα or 20E5. (B) Numbers (cells/mg) of intestinal IL-2Rα^+^ and (C) 20E5^+^ NK cells, in uninfected (uninf) and SE-infected (SE-inf) chickens. Mean + SEM per treatment and time point is shown (*n* = 5).

To determine possible changes in NK cell activation upon SE infection, CD107 surface expression and intracellular IFNγ were analyzed in intestinal and splenic NK cells (Fig 3A). Intestinal NK cells showed a significant increase in surface expression of CD107 and IFNγ production in SE-infected chickens at 1 dpi (11.1±0.1% and 4.6±0.2% respectively) and 3 dpi (9.0±0.4%; 4.1±0.2%) compared to uninfected chickens (Fig 3B and 3C). In the spleen, increased surface expression of CD107 was observed at 1 dpi (8.2±0.5%) and 3 dpi (8.3±0.4%) and increased IFNγ production at 1 dpi (8.2±0.5%), 3 dpi (8.3±0.4%) and 7 dpi (6.4±0.1%) in SE-infected compared to uninfected chickens (Fig 3D and 3E).

**Fig 3.**
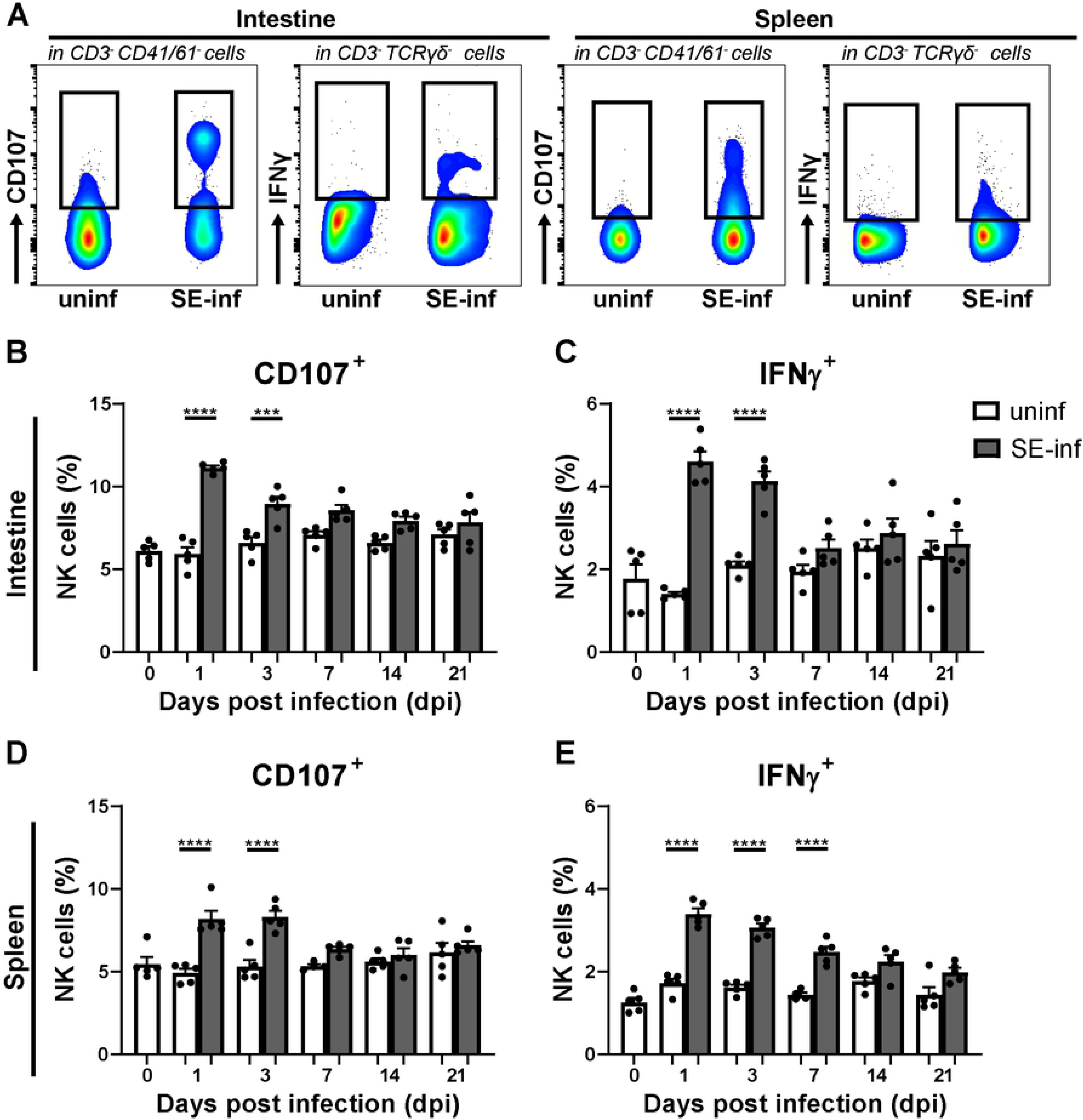
NK cell activation in intestine and spleen of broiler chickens upon SE infection. (A) Gating strategy for NK cells expressing surface CD107 and intracellular IFNγ in the intestine (first and second panels) and spleen (third and fourth panels). (B) Percentages of intestinal NK cells expressing CD107 and (C) IFNγ in uninfected (uninf) and SE-infected (SE-inf) chickens in the course of time. (D) Percentages of splenic NK cells expressing CD107 and (E) IFNγ in uninfected and SE-infected chickens. Mean + SEM per treatment and time point is shown (*n* = 5), for uninfected chickens at 7 dpi in spleen *n* = 4. Statistical significance is indicated as *** *p* < 0.001 and **** *p* < 0.0001.

### Increased presence of APCs in the spleens of SE-infected chickens

To investigate whether infection with SE affects the composition of the APC population, amongst splenocytes, these were stained for APC surface markers and analyzed by flow cytometry. A t-SNE analysis was used to determine the differences in APCs between uninfected and SE-infected chickens (Fig 4A-4C). This analysis clustered cells that have high similarity and separated cells that are unrelated based on the APC surface markers that were used, leading to an unbiased visualization of all cell populations. Two populations were found overrepresented in the spleen of SE-infected chickens (Fig 4B). By gating for subset 1 and 2 and assessing their expression of APC markers, subset 1 was identified as CD11^+^ MRC1LB^+^ and subset 2 as CD11^+^ MRC1LB^−^ (Fig 4C, first panel). In addition, subset 2 could be further divided into two subsets, distinguished by FSC-A and SSC-A characteristics, that were further analyzed separately (Fig 4C, second panel).

**Fig 4.**
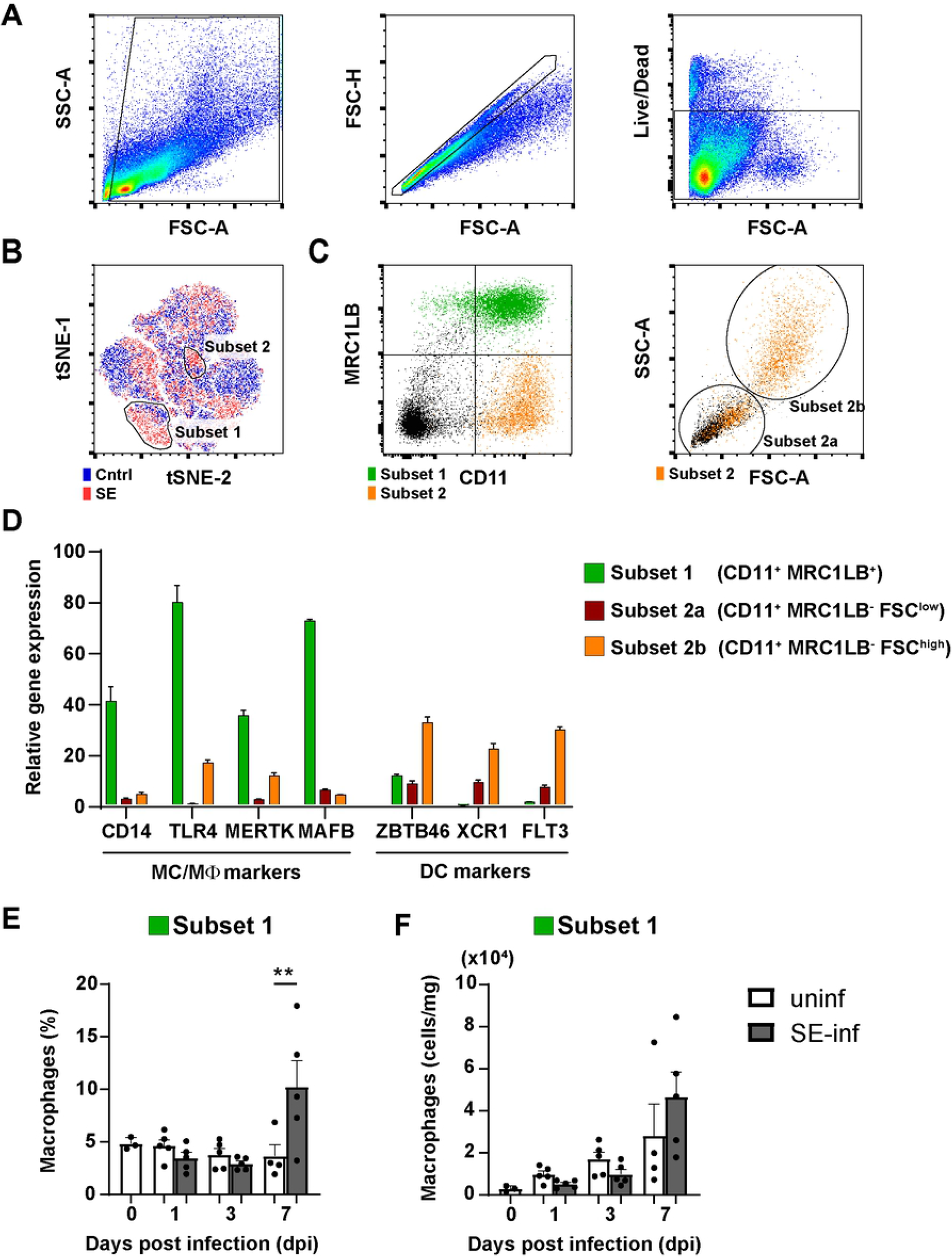
Phenotypic characterization of splenic APCs upon SE infection. (A) Splenocytes were gated for size, excluding debris (FSC-A vs SSC-A), singlets (FSC-A vs FSC-H) and viability (Live/Dead marker-negative) consecutively. (B) A t-SNE analysis was performed on spleen samples of 7 dpi uninfected (uninf, blue) and SE-infected (SE-inf, red) chickens combined. Based on the t-SNE analysis, two population (subset 1 and subset 2) were found enriched among the splenocytes of SE-infected chickens. (C) The populations were evaluated for expression of MRC1LB versus CD11. Subset 2 was evaluated for its FSC-A vs SSC-A scatter pattern and further subdivided into subset 2a and subset 2b. (D) Subset 1 (CD11^+^ MRC1LB^+^), subset 2a (CD11^+^ MRC1LB^−^ FSC^low^) and subset 2b (CD11^+^ MRC1LB^−^ FSC^high^) were sorted by FACS to assess gene expression of immune markers by RT-qPCR relative to the total splenocyte population. (E) The presence and (F) numbers of macrophages in uninfected and SE-infected chickens were assessed over time. Mean + SEM per treatment and time point is shown (*n* = 5), for uninfected chickens at 0 dpi *n* = 3 and at 7 dpi *n* = 4. Statistical significance is indicated as ** *p* < 0.01.

The APC subsets were sorted, and qPCR was performed to compare the expression levels of macrophage- and DC-specific genes between sorted cells and the unsorted total APC population. High expression levels of the monocyte/macrophage genes CD14, TLR4, MERTK and MAFB (Fig 4D) observed on the CD11^+^ MRC1LB^+^ cells, indicates that this subset 1 includes macrophages (hereafter referred to as macrophages). The CD11^+^ MRC1LB^−^ FSC^high^ subset 2b includes DCs as reflected by high expression of the DC genes ZBTB46, XCR1 and FLT3 (hereafter referred to as FSC^high^ DCs) (Fig 4D). The increase in expression of either macrophage- or DC-specific genes was less clear in the CD11^+^ MRC1LB^−^ FSC^low^ subset 2a, however, DC-specific genes were most abundantly expressed (hereafter referred to as FSC^low^ DCs) (Fig 4D).

Next, the percentages (Fig 4E and S2A-B Fig) and numbers (Fig 4F and S2C-D Fig) of the three APC subsets were followed over time in the spleens of uninfected and SE-infected chickens. Due to limited cell numbers, the analysis of APCs could not be performed for two chickens at 0 dpi. At 7 dpi, the percentage of macrophages was significantly increased in SE-infected compared to uninfected chickens (Fig 4E). The FSC^low^ (S3A and S2C Fig) and FSC^high^ (S3B and S2D Fig) DCs were similar in SE-infected compared to uninfected chickens at all time points, although a slight increase in both the percentages and numbers of FSC^high^ DCs was observed at 7 dpi (S2B and S2D Fig).

### APCs become activated in spleens of SE-infected chickens

To assess the activation status of the three APC subsets in response to SE infection, expression levels of immunoglobulin Y receptor CHIR-AB1, co-stimulatory molecules CD40 and CD80, and MHC-II were evaluated by flow cytometry (S3 Fig). The macrophages of SE-infected chickens did not show increased expression of the activation markers compared to uninfected chickens (Fig 5A and 5D and 5G and 5J). Before infection, FSC^low^ DCs of uninfected chickens showed a higher expression of MHC-II (gMFI = 5.0×10^3^±0.2×10^3^ vs 2.4×10^3^±0.1×10^3^, Fig 5K-L respectively) and more cells that were positive for the costimulatory molecules CD40 (27.8±6.5% vs 2.0±0.5%, Fig 5E-F respectively) and CD80 (41.3±3.8% vs 3.8±0.9%, Fig 5H-I respectively) compared to FSC^high^ DCs. At 7 dpi, FSC^low^ DCs showed significantly increased expression of CHIR-AB1 (64.1±5.2% vs 39.9±4.4%, Fig 5B), CD40 (55.1±5.5% vs 30.2±4.3%, Fig 5E) and CD80 (67.7±2.7% vs 42.1±3.9%, Fig 5H) in SE-infected chickens as compared to uninfected chickens. The FSC^high^ DCs showed at 3 dpi trends towards increased expression of CD40 (6.6±1.5% vs 3.7±0.5%, Fig 5F) and CD80 (7.4±1.7% vs 4.0±0.5%, Fig 5I) in SE-infected compared to uninfected chickens, and showed at 7 dpi significantly increased expression of CHIR-AB1 (84.3±5.0% vs 27.5±8.8%, Fig 5C) and MHC-II (gMFI = 4.9×10^3^±0.2×10^3^ vs 3.3×10^3^±0.5×10^3^, Fig 5L) in SE-infected compared to uninfected chickens.

**Fig 5.**
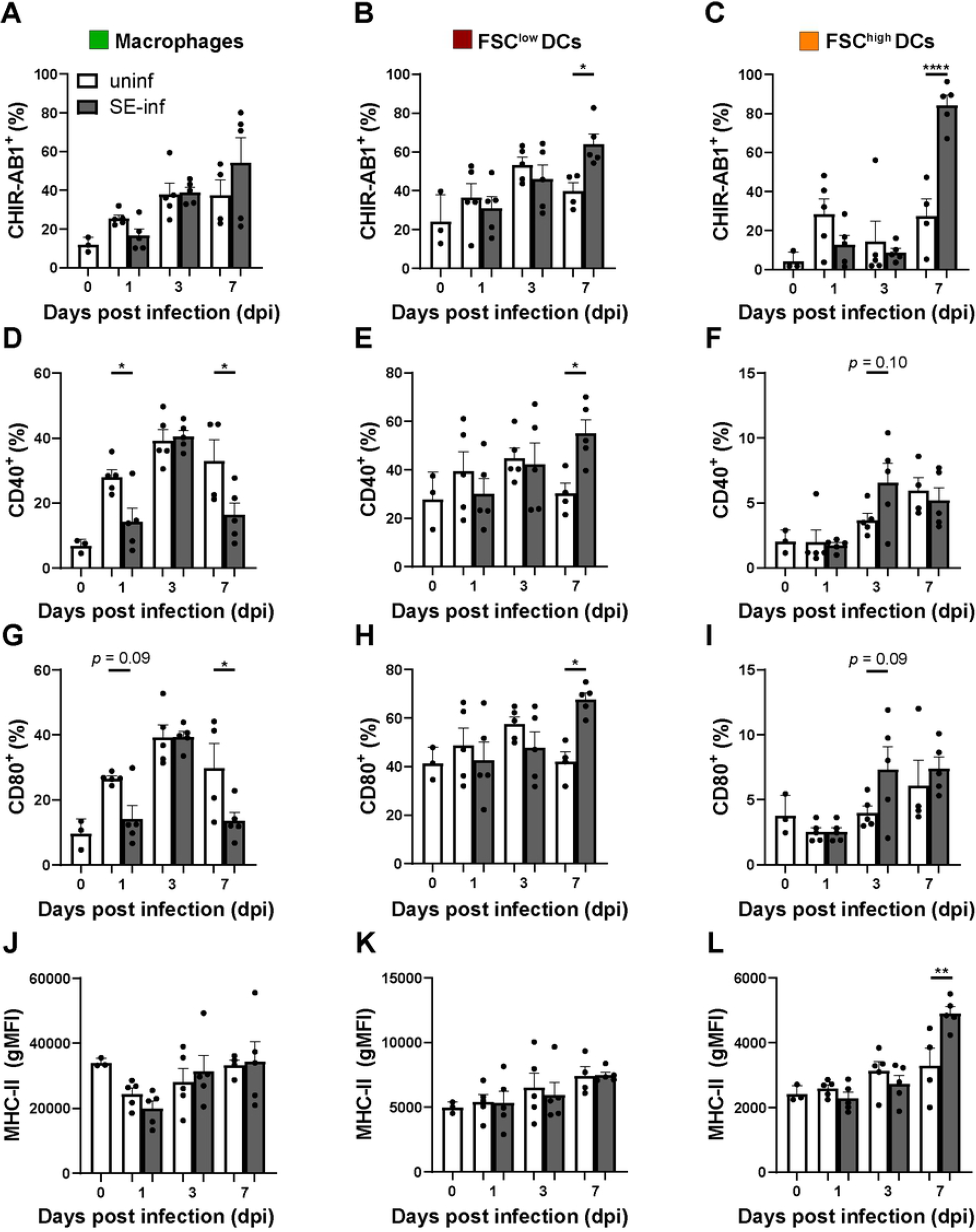
Activation marker expression by splenic APC subsets upon SE infection. (A-C) Macrophages, FSC^low^ DCs and FSC^high^ DCs were assessed over time for CHIR-AB1, (D-F) CD40, (G-I) CD80 and (J-L) MHC-II expression in uninfected (uninf) and SE-infected (SE-inf) chickens. For CHIR-AB1, CD40 and CD80, the percentage of cells in each APC subset expressing the respective markers is shown, and for MHC-II the geometric mean fluorescent intensity (gMFI) of each subset, in accordance with the gating strategy depicted in S4 Fig. Mean + SEM per treatment and time point is shown (*n* = 5), for uninfected chickens at 0 dpi *n* = 3 and at 7 dpi *n =* 4. Statistical significance is indicated as * *p* < 0.05, ** *p* < 0.01, **** *p* < 0.0001.

### Increased presence of intestinal γδ T and cytotoxic T cells at 1 dpi and proliferation of SE-specific splenic T cells ex vivo at 21 dpi

Numbers of γδ T cells and cytotoxic (CD8^+^) αβ T cells were determined in course of time in the intestine and spleen of uninfected and SE-infected chickens (Fig 6A). A trend towards increased numbers of intestinal γδ T cells were found at 1 and 3 dpi, as well as at 21 dpi in SE-infected compared to uninfected chickens (Fig 6B). A significant increase in numbers of intestinal cytotoxic CD8^+^ T cells was observed at 1 dpi (2719±313, Fig 6C), and at 3 dpi and 21 dpi numbers of intestinal cytotoxic CD8^+^ T cells tended to be higher in SE-infected compared to uninfected chickens (Fig 6C). In the spleen, numbers of γδ T cells tended to be decreased at 1 dpi but increased at 3 dpi in SE-infected compared to uninfected chickens (Fig 6D). Numbers of splenic cytotoxic CD8^+^ T cells were similar between uninfected and SE-infected chickens during the course of infection (Fig 6E). Next, γδ T cells and cytotoxic αβ T cells were analyzed for their CD8αα and CD8αβ expression (S4A Fig). Numbers of intestinal CD8αα^+^ γδ T cells (S4B Fig) were significantly enhanced at 1 dpi (1084±133) and 21 dpi (694±240), and CD8αβ^+^ γδ T cell numbers (S4C Fig) tended to increase at those time points in SE-infected compared to uninfected chickens. Similarly, numbers of intestinal cytotoxic CD8αα^+^ T cells (S4D Fig) were significantly enhanced at 1 dpi (1550±242) and tended to be higher at 21 dpi. The cytotoxic CD8αβ^+^ T cell numbers (S4E Fig) showed trends to increase at those time points in SE-infected compared to uninfected chickens. In the spleen, CD8αα^+^ γδ T cell numbers (S4F Fig) were higher at 14 dpi (6537±1476), whereas numbers of CD8αβ^+^ γδ T cells (S4G Fig) were similar in SE-infected versus uninfected chickens. Numbers of splenic cytotoxic CD8αα^+^ (S4H Fig) and CD8αβ^+^ (S4I Fig) T cells as well helper CD4^+^ T cells (S5 Fig) were similar between uninfected and SE-infected chickens during the course of infection. Finally, no significant differences were observed in T cell activation, determined by CD107 and IFNγ expression, in the intestine and spleen between uninfected and SE-infected chickens (S6 Fig). Trends to increase were observed at 3 dpi of CD107 expression by intestinal and splenic CD8^+^ T cells (comprising both γδ and αβ TCRs, S6A-B Fig respectively), and IFNγ production by splenic CD4^+^ T cells (S6D Fig) in SE-infected compared to uninfected chickens. IFNγ production by T cell subsets in the intestine could not be determined due to too low cell numbers.

**Fig 6.**
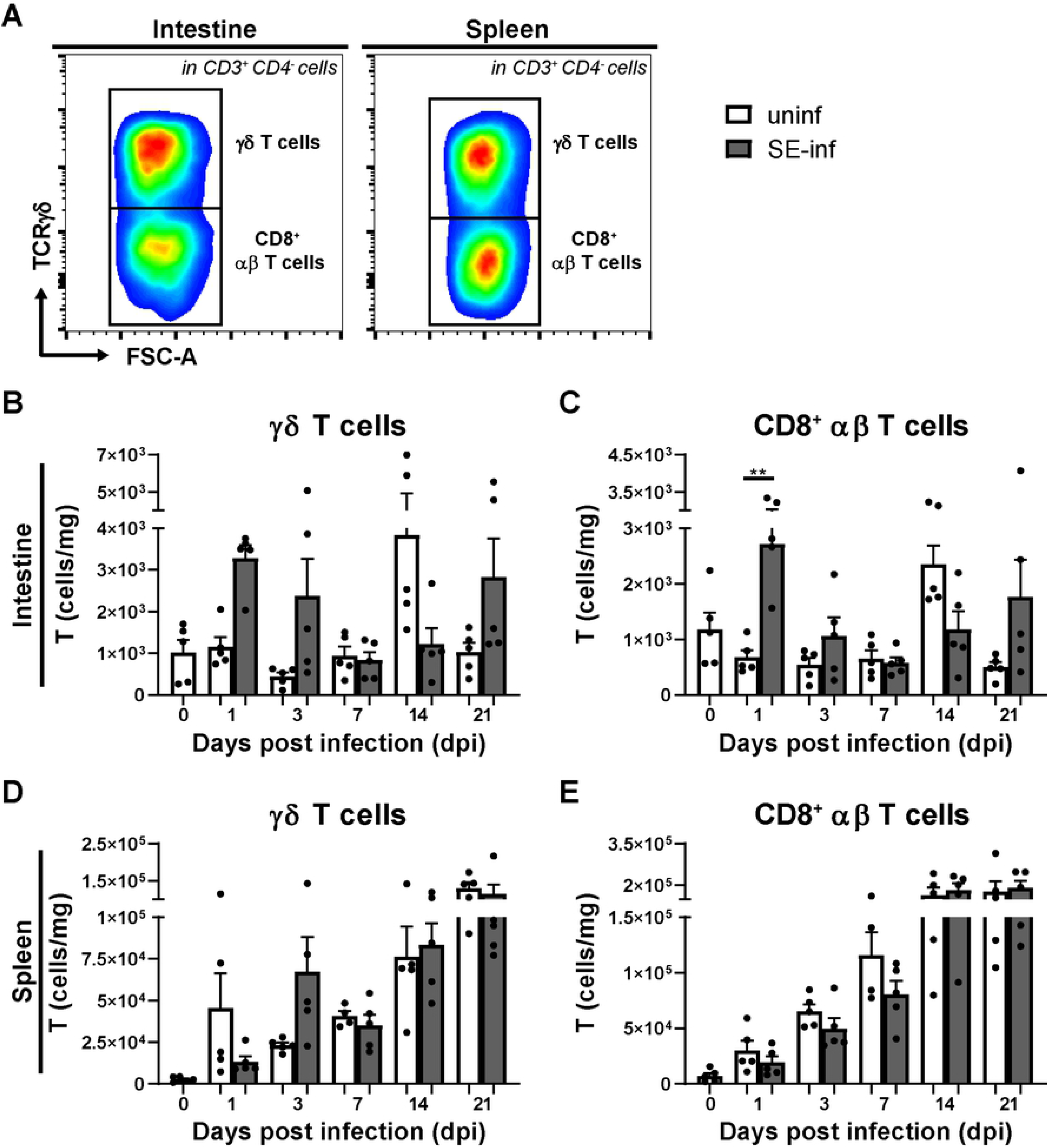
Numbers of intestinal and splenic T cells in broiler chickens upon SE infection. (A) Gating strategy for γδ T cells and CD8^+^ αβ T cells in intestine (first panel) and spleen (second panel). (B) Numbers (cells/mg) of intestinal γδ T cells and (C) CD8^+^ αβ T cells in uninfected (uninf) and SE-infected (SE-inf) chickens. (D) Numbers (cells/mg) of splenic γδ T cells and (E) CD8^+^ αβ T cells in uninfected and SE-infected chickens. Mean + SEM per treatment and time point is shown (*n* = 5), for uninfected chickens at 7 dpi in spleen *n* = 4. Statistical significance is indicated as ** *p* < 0.01.

SE-specific proliferation of T cells, isolated from spleen at 21 dpi, was determined ex vivo (Fig 7A). Increased proliferation of SE-specific CD4^+^ as well as CD8^+^ T cells isolated from SE-infected chickens was observed. This proliferation was antigen dose-dependent, whereas T cells from uninfected chickens did not proliferate upon exposure to inactivated SE (Fig 7B-C).

**Fig 7.**
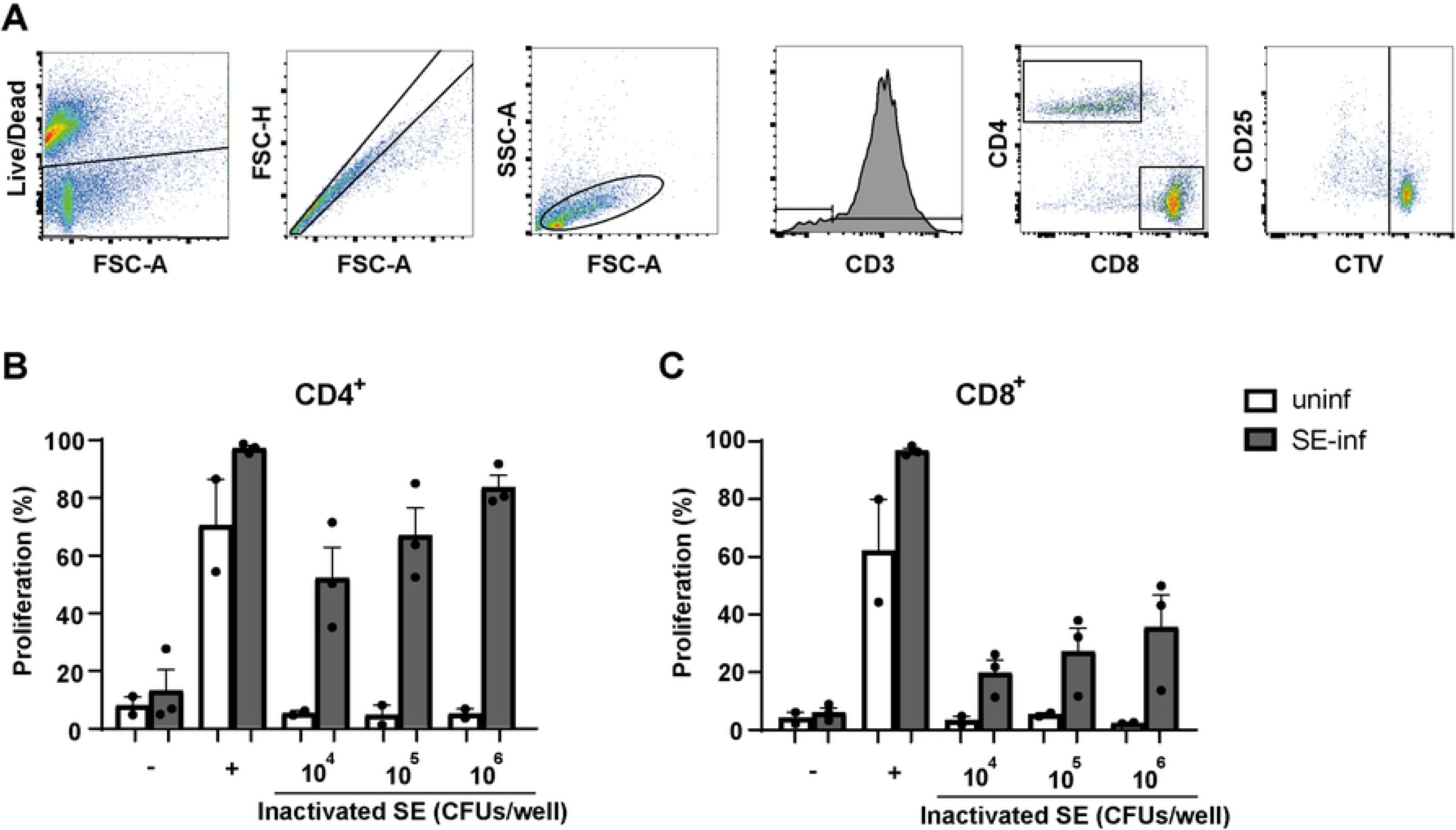
Ex vivo proliferation of SE-specific splenic CD4^+^ and CD8^+^ T cells. (A) The gating strategy shows the consecutive selection for viable cells (Live/Dead marker-negative), single cells (FSC-A vs FSC-H), lymphocytes (FSC-A vs SSC-A) and CD3^+^ T cells. T cells were subdivided into CD4^+^ and CD8^+^ T cells. The final gating step selects on T cell subsets which have divided at least once based on dilution of the cell proliferation dye CellTrace Violet (CTV). (B) The percentage of cells that have proliferated is shown for splenic CD4^+^ and (C) CD8^+^ T cells after four days of stimulation with different doses of formaldehyde-inactivated SE expressed in CFU/well, none stimulated controls (−), or after stimulation with anti-CD3, anti-CD28 and recombinant chicken IL-2 (+). All splenocyte samples were stimulated and measured in triplicate for each of the conditions. Mean + SEM is shown; *n* = 2 for uninfected (uninf) and *n* = 3 for SE-infected (SE-inf) chickens.

### High SE-specific antibody responses were found in all SE-infected chickens after three weeks of infection

The presence of SE-specific antibodies was determined in serum before and after infection in uninfected and SE-infected chickens. In SE-infected chickens, SE-specific antibodies were first detected at 7 dpi, when two out of five chickens showed low antibody titers, that increased in course of time (Fig 8). At 14 dpi SE-specific antibodies were observed in all SE-infected chickens although two chickens showed only low titers. At 21 dpi all SE-infected chickens showed high SE-specific antibody responses with average titers of 455±133. SE-specific antibodies were not found in sera of the uninfected chickens (Fig 8).

**Fig 8.**
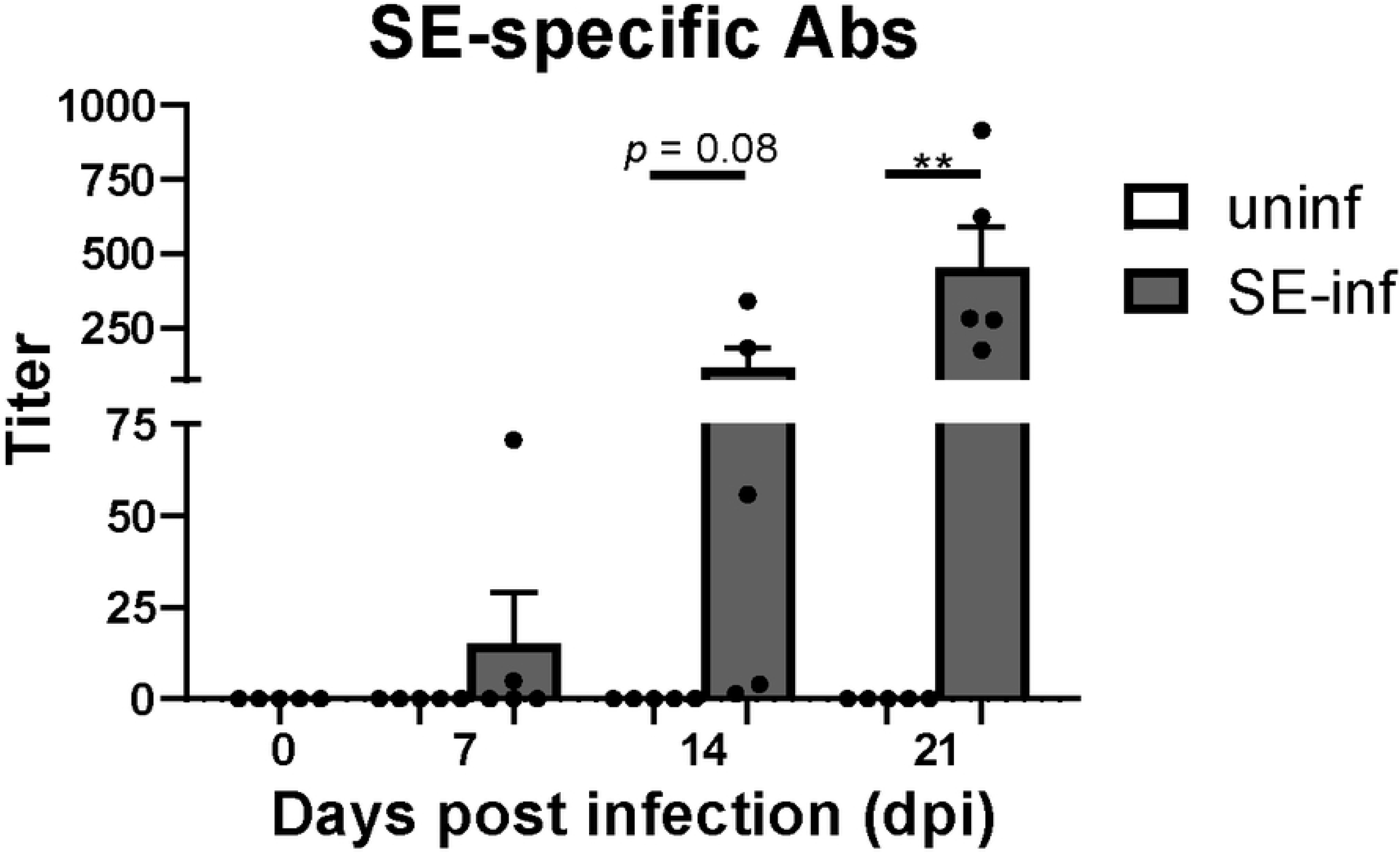
Serum antibody titers in broiler chickens as a response to SE infection. Titers of SE-specific antibodies in sera of uninfected (uninf) and SE-infected (SE-inf) chickens in course of time. Mean + SEM per treatment and time point is shown (*n* = 5) and statistical significance is indicated as ** *p* < 0.01.

## Discussion

In the current study we aimed to provide a detailed analysis of SE-related innate and adaptive immune responses in young broiler chickens up to four weeks of age, to better understand how the immune response contributes to the elimination of infection in course of time. For this purpose, the presence and function of NK cells, various types of APCs and T cells in ileum, as the present infection site and in spleen, as indication of systemic dissemination of SE, were investigated, as well as SE-specific antibody responses in serum, another systemic dissemination indicator. Seven-day-old broiler chickens were successfully infected with SE as was demonstrated by the presence of bacteria in ileum and spleen and the detection of SE-specific T cell proliferation and antibodies. From 1 dpi onwards, increased numbers and activation of innate immune cells as well as numbers of T cells were observed in the intestine and spleen of SE-infected compared to uninfected chickens. From 21 dpi onwards, enhanced T cell numbers in intestine and antibody titers in serum were observed in SE-infected compared to uninfected chickens, resulting in the reduction of SE.

Although SE was demonstrated in ileum and spleen of infected chickens, the presence of the bacteria did not affect growth nor induced severe disease symptoms similar to previous studies in layer chickens [13,34] and young broiler chickens [32,52]. The absence of severe disease symptoms is related to the SE-dose, which was chosen to avoid welfare issues in the chickens. Other studies reported the presence of SE for a longer period in the caecum [32,33], suggesting that SE may prefer colonization in the caecum rather than the ileum.

The enhanced activation of intestinal and splenic NK cells upon SE infection, represented by enhanced CD107 expression and IFNγ production, is in agreement with other studies in chickens showing upregulated mRNA levels of intestinal IFNγ [13] and cytotoxicity-related NK cell genes [12] in young chickens. Our observations are also supported by studies in humans and mice, which reported increased cytotoxicity [14], IFNγ production [16] and CD107 expression [15] of intestinal and systemic NK cells after *Salmonella enterica* serotype Typhimurium infection. The enhanced NK cell activation paralleled increased numbers of intestinal IL-2Rα^+^ and 20E5^+^ NK cells. Due to incompatibility of available reagents, we were not able to determine CD107 and IFNγ expression within the IL-2Rα^+^ and 20E5^+^ NK cell subsets. For that reason, we sorted IL-2Rα^+^ and 20E5^+^ NK cells to perform RT-qPCR and both NK cell subsets showed mRNA levels of *NFIL3*, *IL-7Rα* and *PRF1* genes albeit to different degrees, suggesting that both may be implicated in cytokine production [41–43] as well as cytotoxic activity [43,46] in response to SE infection. The observation that both NK cell subsets are involved in cytotoxicity is confirmed by previous studies in chickens, in which both NK cell subsets in the intestine and spleen showed CD107 expression [7] and intestinal IL-2Rα^+^ NK cells exerted cytotoxicity [8]. In humans, the peripheral IL-2Rα^+^ NK cell population expanded and increased their cytotoxic activity after Toll-like receptor (TLR) stimulation [53].

As SE might activate NK cells directly through TLRs [53–55], more intestinal NK cells as well as enhanced activation, such as cytotoxicity and IFNγ production, may increase the resistance of chickens against SE infection. IFNγ has been reported to activate macrophages resulting in improved clearance of engulfed bacteria [56] and enhanced antigen presentation by APCs inducing T cell responses [57]. Although NK cell activation did not reduce SE numbers in the first week after infection, they may play an important role in the resistance to SE infection by influencing both the innate and adaptive immune cells.

The systemic spread of SE infection as observed, coincided with an increased presence of CD11^+^ MRC1LB^+^ macrophages in the spleen at 7 dpi when bacterial counts were highest. Previous studies have shown that these CD11^+^ MRC1LB^+^ macrophages are largely present in peri-ellipsoid lymphocyte sheaths of the spleen [58], and have a role in clearing blood-borne bacteria in chickens [59], equivalent to that of the marginal zone macrophages in mammals [60]. Therefore, these macrophages are suggested to be involved in clearing the SE from 7 dpi onwards. The FSC^high^ DCs tended to be present in increasing numbers and showed increased expression of CHIR-AB1 and MHC-II at 7 dpi in SE-infected chickens compared to uninfected chickens, indicating a role in antigen presentation. The FSC^low^ DCs did not increase in numbers but showed an increased expression of the activation markers CHIR-AB1, CD40 and CD80 in SE-infected chickens. The two DC subsets were highly similar, and might comprise DCs at different stages of maturation with the FSC^low^ subset being more mature based on the expression of CD40, CD80 and MHCII [22,49,61]. These results suggests that the increased presence of macrophages clear bacteria initially and the increased activation of DC subsets contribute to antigen presentation to stimulate the adaptive immune responses, all together resulting in further reduction of SE in infected chickens.

Whereas the APC subsets are likely to contribute to the clearance of the bacteria, it has also been suggested that they may worsen the impact of infection by acting as a carrier for *Salmonellae* [23] and contribute to systemic dissemination [28], since this bacterium is able to survive intracellularly in chicken macrophages [26,27] and DCs [61]. The ability of *Salmonellae* to inhibit activation of APCs might explain why NK cells showed earlier activation than APCs and the high presence of SE found at 7 dpi in our study. Other studies have demonstrated that *Salmonella*-infected APCs secrete IL-12/IL-18 resulting in enhanced expression of the early activation marker IL-2Rα^+^ on NK cells, thereby inducing their activation by increased cytotoxicity and IFNγ production [15]. This IFNγ production can subsequently stimulate additional macrophages to clear phagocytosed bacteria [62] or impair intracellular survival by direct killing of infected-macrophages [15], which might be involved in the reduction of SE towards and after 7 dpi observed in our study.

T cell presence and SE-specific T cell proliferation and antibodies were addressed as well. All infected chickens in our study had circulating antibodies specific for SE after three weeks, which was in agreement with other studies [32,33]. The observed T cell responses are similar to increased numbers of intestinal and splenic γδ and cytotoxic αβ T cells in response to SE early and three weeks after infection [13,17,18], as well as to increased CD8αα^+^ γδ T cell numbers [29,63] in prior studies in chickens. The increase in cytotoxic CD8αα^+^ T cell numbers, however, has not been shown before in chickens in response to SE infection. The more innate-like nature of γδ T cell responses early after infection have been recognized as well as the antigen-specific responses of cytotoxic CD8^+^ T cells approximately two weeks after infection, whereas the functional difference between CD8αα and CD8αβ expression is less clear [64–67]. Although expression of CD107 and IFNγ production by intestinal and splenic γδ T cells, cytotoxic CD8^+^ T cells, and splenic helper CD4^+^ T cells did not significantly differ between uninfected and SE-infected chickens, proliferation of SE-specific splenic T cells of SE-infected chickens ex vivo was observed three weeks after infection and not in uninfected chickens. The initial increased presence of T cells in our study did not result in reduction of SE, however, the SE-specific T cells and antibodies in course of infection are suggested to reduce the number of SE.

We propose the following model of SE-induced immune responses. At the start of a SE infection, NK cells become activated either directly through TLRs or indirectly by interaction with activated macrophages that have engulfed SE through contact and cytokine production. Moreover, intestinal γδ T cells and cytotoxic CD8^+^ T cells are involved in the first response controlling SE infection by cytokine production. Subsequently, NK cells kill infected cells directly by degranulation and indirectly by IFNγ production, which in turn stimulates macrophages to destroy engulfed bacteria. The destruction of bacteria by APCs results in antigens that can be presented to SE-specific CD4^+^ and CD8^+^ T cells, which differentiate into CD4^+^ helper T cells and CD8^+^ cytotoxic T cells, respectively. The CD4^+^ T helper cells stimulate the differentiation of SE-specific B cells into antibody-producing plasma cells and secrete cytokines leading to further stimulation of NK cells, APCs and cytotoxic CD8^+^ T cells, thereby creating a positive feedback loop, resulting in the reduction of SE and thereby elimination of the infection.

In conclusion, this study shows that *Salmonella enterica* serotype Enteritidis infection in young broiler chickens firstly increases local and systemic presence and activation of NK cells as well as local presence of T cells, followed by increased presence of macrophages and activation of DCs. Subsequently, specific T cell and antibody responses are induced, all together resulting in reduction of SE. These insights in understanding the response of innate and adaptive immune cells upon SE infection will aid in developing immune-modulation strategies to stimulate innate cells. The potential strengthening of immune responsiveness by vaccines or feed strategies during early life may increase resistance and may prevent SE infection and colonization in young broiler chickens, eventually increasing the food safety for humans.

## Acknowledgements

We thank the animal caretakers and F.C. Velkers of the Department Population Health Sciences, division Farm Animal Health, Faculty of Veterinary Medicine, Utrecht University, for their help during the animal experiments and SE inoculation. We acknowledge E. Broens and A.J. Timmerman of the VMDC, Faculty of Veterinary Medicine, Utrecht University, for kindly providing the SE strain and help with the SE culture. We are thankful to dr. I.S. Ludwig for her help during the isolation of immune cells and we acknowledge dr. G.J.A. Arkesteijn for maintaining optimal working conditions of the Flow Cytometry and Cell Sorting Facility, Faculty of Veterinary Medicine, Utrecht University.

## Supporting Information Captions

**S1 Fig. Effect of SE infection on numbers of splenic NK cells in broiler chickens.** (A) Gating strategy for CD3 negative cells expressing either IL-2Rα or 20E5. (B) Numbers (cells/mg) of splenic IL-2Rα^+^ and (C) 20E5^+^ NK cells in uninfected (uninf) and SE-infected (SE-inf) chickens in the course of time. (D) Gene expression levels of NK cell lineage marker (NFIL3), IL-7Rα and perforin 1 (PRF1) by RT-qPCR in sorted IL-2Rα^+^ and 20E5^+^ NK cell subsets. Mean + SEM per treatment and time point is shown (*n* = 5), for uninfected chickens at 7 dpi *n* = 4 and for gene expression levels *n* = 1.

**S2 Fig. Phenotypic characterization of splenic APCs upon SE infection.** (A-B) The presence and (C-D) numbers of FSC^low^ DCs and and FSC^high^ DCs in uninfected (uninf) and SE-infected (SE-inf) chickens were assessed over time. Mean + SEM per treatment and time point is shown (*n =* 5), for uninfected chickens at 0 dpi *n* = 3 and at 7 dpi *n* = 4. Statistical significance is indicated as ** *p* < 0.01.

**S3 Fig. The gating strategy used to determine the activation status of the APC subsets as depicted in Fig 5.** The three identified splenic APC subsets (A) macrophages, (B) FSC^low^ DCs and (C) FSC^high^ DCs were assessed for CHIR-AB1, CD40, CD80 and MHC-II. For CHIR-AB1, CD40 and CD80, the cells expressing the respective markers were selected and expressed as a percentage. The expression of MHC-II by each subset was expressed as the geometric mean fluorescent intensity (gMFI).

**S4 Fig. Numbers of intestinal and splenic γδ T cells and cytotoxic T cells expressing either CD8αα or CD8αβ in broiler chickens upon SE infection.** (A) Gating strategy for γδ T cells and CD8^+^ αβ T cells expressing either CD8αα or CD8αβ in intestine (first and second panels) and spleen (third and fourth panels). (B) Numbers (cells/mg) of intestinal CD8αα^+^ γδ T cells, (C) CD8αβ^+^ γδ T cells, (D) cytotoxic CD8αα^+^ T cells and (E) CD8αβ^+^ T cells in uninfected (uninf) and SE-infected (SE-inf) chickens in the course of time. (F) Numbers (cells/mg) of splenic CD8αα^+^ γδ T cells, (G) CD8αβ^+^ γδ T cells, (H) cytotoxic CD8αα^+^ T cells and (I) CD8αβ^+^ T cells in uninfected and SE-infected chickens. Mean + SEM per treatment and time point is shown (*n* = 5), for uninfected chickens at 1 dpi in intestine and spleen *n* = 4 due to numbers of events acquired in the gate of interest were < 100, and at 7 dpi in spleen *n* = 4. Statistical significance is indicated as * *p* < 0.05, ** *p* < 0.01. *** *p* < 0.001.

**S5 Fig. Numbers of CD4^+^ T cells in the spleen of broiler chickens upon SE infection.** Numbers (cells/mg) of splenic CD4^+^ αβ T cells in uninfected (uninf) and SE-infected (SE-inf) chickens in the course of time. Mean + SEM per treatment and time point is shown (*n* = 5), for uninfected chickens at 7 dpi *n* = 4.

**S6 Fig. T cell activation in the intestine and spleen of broiler chickens upon SE infection.** (A) Percentages of intestinal CD8^+^ T cells expressing CD107 (including both γδ and αβ T cells) in uninfected (uninf) and SE-infected (SE-inf) chickens in the course of time. (B) Percentages of splenic CD8^+^ T cells expressing CD107 (including both γδ and αβ T cells), (C) CD8^+^ γδ T cells expressing IFNγ, (D) CD4^+^ αβ T cells expressing IFNγ and (E) CD8^+^ αβ T cells expressing IFNγ in uninfected (uninf) and SE-infected (SE-inf) chickens over time. Mean + SEM per treatment and time point is shown (*n* = 5), for uninfected chickens at 7 dpi in spleen *n* = 4 and at 1 and 3 dpi in intestine percentages were not determined (n.d.) due to numbers of events acquired in the gate of interest were < 100.

## References

1. Kallapura G, Morgan MJ, Pumford NR, Bielke LR, Wolfenden AD, Faulkner OB, et al. Evaluation of the respiratory route as a viable portal of entry for Salmonella in poultry via intratracheal challenge of Salmonella Enteritidis and Salmonella Typhimurium. Poult Sci. 2014;93: 340–346. doi: 10.3382/ps.2013-03602.

2. Kallapura G, Kogut MH, Morgan MJ, Pumford NR, Bielke LR, Wolfenden AD, et al. Fate of Salmonella Senftenberg in broiler chickens evaluated by challenge experiments. Avian Pathol. 2014;43: 305–309. doi: 10.1080/03079457.2014.923554.

3. Suzuki S. Pathogenicity of Salmonella enteritidis in poultry. Int J Food Microbiol. 1994;21: 89–105. doi: 10.1016/0168-1605(94)90203-8.

4. World Organisation for Animal Health, (OIE). Prevention, detection and control of Salmonella in poultry. In: OIE, editor. Terrestrial Animal Health Code. Paris: Office International des Epizooties; 2019.

5. Sharma JM, Tizard I. Avian cellular immune effector mechanisms - a review. Avian Pathol. 1984;13: 357–376. doi: 10.1080/03079458408418541.

6. Klasing KC LT. Functions, costs and benefits of the immune system during development and growth. . 1999;In: Adams NJ, and Slotow R (eds): 2817–2835.

7. Meijerink N, van Haarlem DA, Velkers FC, Stegeman AJ, Rutten, V. P. M. G., Jansen CA. Analysis of chicken intestinal natural killer cells, a major IEL subset during embryonic and early life. Dev Comp Immunol. 2020: 103857. doi: S0145-305X(20)30412-2 [pii].

8. Göbel TWF, Kaspers B, Stangassinger M. NK and T cells constitute two major, functionally distinct intestinal epithelial lymphocyte subsets in the chicken. Int Immunol. 2001;13: 757–762. doi: 10.1093/intimm/13.6.757.

9. Fenzl L, Göbel TW, Neulen M-. γδ T cells represent a major spontaneously cytotoxic cell population in the chicken. Dev Comp Immunol. 2017;73: 175–183. doi: 10.1016/j.dci.2017.03.028.

10. Lillehoj HS, Trout JM. Avian gut-associated lymphoid tissues and intestinal immune responses to Eimeria parasites. Clin Microbiol Rev. 1996;9: 349–360. doi: 10.1128/cmr.9.3.349.

11. Schokker D, De Koning D-, Rebel JMJ, Smits MA. Shift in chicken intestinal gene association networks after infection with Salmonella. Comp Biochem Physiol Part D Genomics Proteomics. 2011;6: 339–347. doi: 10.1016/j.cbd.2011.07.004.

12. Schokker D, Smits MA, Hoekman AJW, Parmentier HK, Rebel JMJ. Effects of Salmonella on spatial-temporal processes of jejunal development in chickens. Dev Comp Immunol. 2010;34: 1090–1100. doi: 10.1016/j.dci.2010.05.013.

13. Carvajal BG, Methner U, Pieper J, Berndt A. Effects of Salmonella enterica serovar Enteritidis on cellular recruitment and cytokine gene expression in caecum of vaccinated chickens. Vaccine. 2008;26: 5423–5433. doi: 10.1016/j.vaccine.2008.07.088.

14. Schafer R, Eisenstein TK. Natural killer cells mediate protection induced by a Salmonella aroA mutant. Infect Immun. 1992;60: 791–797.

15. Lapaque N, Walzer T, Méresse S, Vivier E, Trowsdale J. Interactions between human NK cells and macrophages in response to Salmonella infection. J Immunol. 2009;182: 4339–4348. doi: 10.4049/jimmunol.0803329.

16. Harrington L, Srikanth CV, Antony R, Shi HN, Cherayil BJ. A role for natural killer cells in intestinal inflammation caused by infection with Salmonella enterica serovar Typhimurium. FEMS Immunol Med Microbiol. 2007;51: 372–380. doi: 10.1111/j.1574-695X.2007.00313.x.

17. Schokker D, Peters THF, Hoekman AJW, Rebel JMJ, Smits MA. Differences in the early response of hatchlings of different chicken breeding lines to Salmonella enterica serovar Enteritidis infection. Poult Sci. 2012;91: 346–353. doi: 10.3382/ps.2011-01758.

18. Van Hemert S, Hoekman AJW, Smits MA, Rebel JMJ. Immunological and gene expression responses to a Salmonella infection in the chicken intestine. Vet Res. 2007;38: 51–63. doi: 10.1051/vetres:2006048.

19. van Hemert S, Hoekman AJW, Smits MA, Rebel JMJ. Gene expression responses to a Salmonella infection in the chicken intestine differ between lines. Vet Immunol Immunopathol. 2006;114: 247–258. doi: 10.1016/j.vetimm.2006.08.007.

20. van Hemert S, Hoekman AJW, Smits MA, Rebel JMJ. Early host gene expression responses to a Salmonella infection in the intestine of chickens with different genetic background examined with cDNA and oligonucleotide microarrays. Comp Biochem Physiol Part D Genomics Proteomics. 2006;1: 292–299. doi: 10.1016/j.cbd.2006.05.001.

21. Staines K, Hunt LG, Young JR, Butter C. Evolution of an expanded mannose receptor gene family. PLoS ONE. 2014;9. doi: 10.1371/journal.pone.0110330.

22. Kamble NM, Jawale CV, Lee JH. Activation of chicken bone marrow-derived dendritic cells induced by a Salmonella Enteritidis ghost vaccine candidate. Poult Sci. 2016;95: 2274–2280. doi: 10.3382/ps/pew158.

23. Chappell L, Kaiser P, Barrow P, Jones MA, Johnston C, Wigley P. The immunobiology of avian systemic salmonellosis. Vet Immunol Immunopathol. 2009;128: 53–59. doi: 10.1016/j.vetimm.2008.10.295.

24. Gomes AVS, Quinteiro-Filho WM, Ribeiro A, Ferraz-de-Paula V, Pinheiro ML, Baskeville E, et al. Overcrowding stress decreases macrophage activity and increases Salmonella Enteritidis invasion in broiler chickens. Avian Pathol. 2014;43: 82–90. doi: 10.1080/03079457.2013.874006.

25. Vazquez-Torres A, Xu Y, Jones-Carson J, Holden DW, Lucia SM, Dinauer MC, et al. Salmonella pathogenicity island 2-dependent evasion of the phagocyte NADPH oxidase. Science. 2000;287: 1655–1658. doi: 10.1126/science.287.5458.1655.

26. Jones MA, Wigley P, Page KL, Hulme SD, Barrow PA. Salmonella enterica serovar Gallinarum requires the Salmonella pathogenicity island 2 type III secretion system but not the Salmonella pathogenicity island 1 type III secretion system for virulence in chickens. Infect Immun. 2001;69: 5471–5476. doi: 10.1128/IAI.69.9.5471-5476.2001.

27. Jones MA, Hulme SD, Barrow PA, Wigley P. The Salmonella pathogenicity island 1 and Salmonella pathogenicity island 2 type III secretion systems play a major role in pathogenesis of systemic disease and gastrointestinal tract colonization of Salmonella enterica serovar Typhimurium in the chicken. Avian Pathol. 2007;36: 199–203. doi: 10.1080/03079450701264118.

28. Vazquez-Terres A, Jones-Carson J, Bäumler AJ, Falkow S, Valdivia R, Brown W, et al. Extraintestinal dissemination of Salmonella by CD18-expressing phagocytes. Nature. 1999;401: 804–808. doi: 10.1038/44593.

29. Berndt A, Pieper J, Methner U. Circulating γδ T cells in response to Salmonella enterica serovar enteritidis exposure in chickens. Infect Immun. 2006;74: 3967–3978. doi: 10.1128/IAI.01128-05.

30. Sekelova Z, Polansky O, Stepanova H, Fedr R, Faldynova M, Rychlik I, et al. Different roles of CD4, CD8 and γδ T-lymphocytes in naive and vaccinated chickens during Salmonella Enteritidis infection. Proteomics. 2017;17. doi: 10.1002/pmic.201700073.

31. Desmidt M, Ducatelle R, Mast J, Goddeeris BM, Kaspers B, Haesebrouck F. Role of the humoral immune system in Salmonella enteritidis phage type four infection in chickens. Vet Immunol Immunopathol. 1998;63: 355–367. doi: 10.1016/S0165-2427(98)00112-3.

32. Zhen W, Shao Y, Gong X, Wu Y, Geng Y, Wang Z, et al. Effect of dietary Bacillus coagulans supplementation on growth performance and immune responses of broiler chickens challenged by Salmonella enteritidis. Poult Sci. 2018;97: 2654–2666. doi: 10.3382/ps/pey119.

33. Berthelot-Hérault F, Mompart F, Zygmunt MS, Dubray GE, Duchet-Suchaux M. Antibody responses in the serum and gut of chicken lines differing in cecal carriage of Salmonella enteritidis. Vet Immunol Immunopathol. 2003;96: 43–52. doi: 10.1016/S0165-2427(03)00155-7.

34. Van De Reep L, Nielen M, Verstappen, K. M. H. W., Broens EM, Van Den Broek J, Velkers FC. Response to a Salmonella Enteritidis challenge in old laying hens with different vaccination histories. Poult Sci. 2018;97: 2733–2739. doi: 10.3382/ps/pey134.

35. Meijerink N, Kers JG, Velkers FC, van Haarlem DA, Lamot DM, de Oliveira JE, et al. Early Life Inoculation With Adult-Derived Microbiota Accelerates Maturation of Intestinal Microbiota and Enhances NK Cell Activation in Broiler Chickens. Front Vet Sci. 2020;7. doi: 10.3389/fvets.2020.584561.

36. Jansen CA, van de Haar, P. M., van Haarlem D, van Kooten P, de Wit S, van Eden W, et al. Identification of new populations of chicken natural killer (NK) cells. Dev Comp Immunol. 2010;34: 759–767. doi: 10.1016/j.dci.2010.02.009.

37. Ariaans MP, van de Haar, P. M., Lowenthal JW, van Eden W, Hensen EJ, Vervelde L. ELISPOT and intracellular cytokine staining: Novel assays for quantifying T cell responses in the chicken. Dev Comp Immunol. 2008;32: 1398–1404. doi: 10.1016/j.dci.2008.05.007.

38. Belkina AC, Ciccolella CO, Anno R, Halpert R, Spidlen J, Snyder-Cappione JE. Automated optimized parameters for T-distributed stochastic neighbor embedding improve visualization and analysis of large datasets. Nat Commun. 2019;10. doi: 10.1038/s41467-019-13055-y.

39. van den Biggelaar, R. H. G. A., van Eden W, Rutten, V. P. M. G., Jansen CA. Nitric oxide production and fc receptor-mediated phagocytosis as functional readouts of macrophage activity upon stimulation with inactivated poultry vaccines in vitro. Vaccines. 2020;8: 1–16. doi: 10.3390/vaccines8020332.

40. Eldaghayes I, Rothwell L, Williams A, Withers D, Balu S, Davison F, et al. Infectious bursal disease virus: Strains that differ in virulence differentially modulate the innate immune response to infection in the chicken bursa. Viral Immunol. 2006;19: 83–91. doi: 10.1089/vim.2006.19.83.

41. Michaud A, Dardari R, Charrier E, Cordeiro P, Herblot S, Duval M. IL-7 enhances survival of human CD56bright NK cells. J Immunother. 2010;33: 382–390. doi: 10.1097/CJI.0b013e3181cd872d.

42. Male V, Nisoli I, Kostrzewski T, Allan DS, Carlyle JR, Lord GM, et al. The transcription factor E4bp4/Nfil3 controls commitment to the NK lineage and directly regulates Eomes and Id2 expression. J Exp Med. 2014;211: 635–642. doi: 10.1084/jem.20132398.

43. Wendt K, Wilk E, Buyny S, Buer J, Schmidt RE, Jacobs R. Gene and protein characteristics reflect functional diversity of CD56 dim and CD56bright NK cells. J Leukocyte Biol. 2006;80: 1529–1541. doi: 10.1189/jlb.0306191.

44. Di Santo JP. A defining factor for natural killer cell development. Nat Immunol. 2009;10: 1051–1052. doi: 10.1038/ni1009-1051.

45. Luevano M, Madrigal A, Saudemont A. Transcription factors involved in the regulation of natural killer cell development and function: An update. Front Immunol. 2012;3. doi: 10.3389/fimmu.2012.00319.

46. Sarson AJ, Abdul-Careem MF, Read LR, Brisbin JT, Sharif S. Expression of cytotoxicity-associated genes in Marek’s disease virus-infected chickens. Viral Immunol. 2008;21: 267–272. doi: 10.1089/vim.2007.0094.

47. Livak KJ, Schmittgen TD. Analysis of relative gene expression data using real-time quantitative PCR and the 2-ΔΔCT method. Methods. 2001;25: 402–408. doi: 10.1006/meth.2001.1262.

48. Manh T-V, Marty H, Sibille P, Vern YL, Kaspers B, Dalod M, et al. Existence of conventional dendritic cells in gallus gallus revealed by comparative gene expression profiling. J Immunol. 2014;192: 4510–4517. doi: 10.4049/jimmunol.1303405.

49. van den Biggelaar, R. H. G. A., Arkesteijn GJA, Rutten, V. P. M. G., van Eden W, Jansen CA. In vitro Chicken Bone Marrow-Derived Dendritic Cells Comprise Subsets at Different States of Maturation. Front Immunol. 2020;11. doi: 10.3389/fimmu.2020.00141.

50. van Haarlem DA, van Kooten, P. J. S., Rothwell L, Kaiser P, Vervelde L. Characterisation and expression analysis of the chicken interleukin-7 receptor alpha chain. Dev Comp Immunol. 2009;33: 1018–1026. doi: 10.1016/j.dci.2009.05.001.

51. Sundick RS, Gill-Dixon C. A Cloned Chicken Lymphokine Homologous to Both Mammalian IL-2 and IL-15. J Immunol. 1997;159: 720–725.

52. Raehtz S, Hargis BM, Kuttappan VA, Pamukcu R, Bielke LR, McCabe LR. High molecular weight polymer promotes bone health and prevents bone loss under salmonella challenge in broiler chickens. Front Physiol. 2018;9. doi: 10.3389/fphys.2018.00384.

53. Rudnicka K, Matusiak A, Chmiela M. CD25 (IL-2R) expression correlates with the target cell induced cytotoxic activity and cytokine secretion in human natural killer cells. Acta Biochim Pol. 2015;62: 885–894. doi: 10.18388/abp.2015_1152.

54. Adib-Conquy M, Scott-Algara D, Cavaillon J-, Souza-Fonseca-Guimaraes F. TLR-mediated activation of NK cells and their role in bacterial/viral immune responses in mammals. Immunol Cell Biol. 2014;92: 256–262. doi: 10.1038/icb.2013.99.

55. Marcenaro E, Ferranti B, Falco M, Moretta L, Moretta A. Human NK cells directly recognize Mycobacterium bovis via TLR2 and acquire the ability to kill monocyte-derived DC. Int Immunol. 2008;20: 1155–1167. doi: 10.1093/intimm/dxn073.

56. Okamura M, Lillehoj HS, Raybourne RB, Babu US, Heckert RA, Tani H, et al. Differential responses of macrophages to Salmonella enterica serovars Enteritidis and Typhimurium. Vet Immunol Immunopathol. 2005;107: 327–335. doi: 10.1016/j.vetimm.2005.05.009.

57. Pallmer K, Oxenius A. Recognition and regulation of T cells by NK cells. Front Immunol. 2016;7. doi: 10.3389/fimmu.2016.00251.

58. Hu T, Wu Z, Bush SJ, Freem L, Vervelde L, Summers KM, et al. Characterization of subpopulations of chicken mononuclear phagocytes that express TIM4 and CSF1R. J Immunol. 2019;202: 1186–1199. doi: 10.4049/jimmunol.1800504.

59. Yu K, Gu MJ, Pyung YJ, Song K-, Park TS, Han SH, et al. Characterization of splenic MRC1hiMHCIIlo and MRC1loMHCIIhi cells from the monocyte/macrophage lineage of White Leghorn chickens. Vet Res. 2020;51. doi: 10.1186/s13567-020-00795-9.

60. A-Gonzalez N, Guillen JA, Gallardo G, Diaz M, De La Rosa, J. V., Hernandez IH, et al. The nuclear receptor LXRα controls the functional specialization of splenic macrophages. Nat Immunol. 2013;14: 831–839. doi: 10.1038/ni.2622.

61. Kamble NM, Jawale CV, Lee JH. Interaction of a live attenuated Salmonella Gallinarum vaccine candidate with chicken bone marrow-derived dendritic cells. Avian Pathol. 2016;45: 235–243. doi: 10.1080/03079457.2016.1144919.

62. Rosenberger CM, Brett Finlay B. Macrophages inhibit Salmonella Typhimurium replication through MEK/ERK kinase and phagocyte NADPH oxidase activities. J Biol Chem. 2002;277: 18753–18762. doi: 10.1074/jbc.M110649200.

63. Pieper J, Methner U, Berndt A. Characterization of avian γδ T-cell subsets after Salmonella enterica serovar typhimurium infection of chicks. Infect Immun. 2011;79: 822–829. doi: 10.1128/IAI.00788-10.

64. Gangadharan D, Cheroutre H. The CD8 isoform CD8αα is not a functional homologue of the TCR co-receptor CD8αβ. Curr Opin Immunol. 2004;16: 264–270. doi: 10.1016/j.coi.2004.03.015.

65. Cheroutre H, Lambolez F, Mucida D. The light and dark sides of intestinal intraepithelial lymphocytes. Nat Rev Immunol. 2011;11: 445–456. doi: 10.1038/nri3007.

66. Pasman L, Kasper DL. Building conventions for unconventional lymphocytes. Immunol Rev. 2017;279: 52–62. doi: 10.1111/imr.12576.

67. Leishman AJ, Naidenko OV, Attinger A, Koning F, Lena CJ, Xiong Y, et al. T cell responses modulated through interaction between CD8αα and the nonclassical MHC class I molecule, TL. Science. 2001;294: 1936–1939. doi: 10.1126/science.1063564.

